# Selenoprotein biosynthesis couples ferroptosis defense to disulfidptosis tolerance

**DOI:** 10.64898/2026.11.17.766756

**Authors:** Hariharan Moorthy, Mostafa Ejtehadifar, Gholamreza Fazeli, Florencio Porto Freitas, Zhiyi Chen, Biplab Ghosh, Jingkun Ma, Ragini Singh, Shibo Sun, Mami Sato, Marietta Fabiano, Ulrich Schweizer, Seiryo Ogata, Uladzimir Barayeu, Takaaki Akaike, Felix C. E. Vogel, Lisa Schlicker, Almut Schulze, Hellmut G. Augustin, Ashok Kumar Jayavelu, Hamed Alborzinia, José Pedro Friedmann Angeli

## Abstract

Selenoprotein biosynthesis is a candidate target for inducing ferroptosis through GPX4 loss. Here we show that disrupting this pathway also promotes cystine uptake that supports ferroptosis defense into a source of toxicity. Loss of selenophosphate synthetase 2 (SEPHS2) impairs both GPX4-mediated lipid protection and thioredoxin reductase 1 (TXNRD1)-dependent reduction of cystine to cysteine, a precursor for glutathione synthesis. Impaired cystine reduction promotes NRF2-dependent induction of SLC7A11 (xCT), creating a feed-forward loop that drives disulfide stress that persists during ferroptosis blockade. Glutathione depletion intensifies this vulnerability, producing protein aggregation and cytoskeletal remodeling associated with disulfidptosis. In neuroblastoma, among the cancers with the lowest SLC7A11 expression, SEPHS2 depletion increases disulfide-linked protein aggregation and suppresses orthotopic xenograft growth. Crucially, forced cystine uptake through SLC7A11 overexpression alone was shown to suppress xenograft growth, despite increasing tumor glutathione. These findings place selenoprotein biosynthesis at the crossroads of ferroptosis and disulfidptosis, revealing that the antioxidant benefit of cystine uptake depends on the reducing capacity required to prevent its toxicity.

## Introduction

The selenoprotein glutathione peroxidase 4 (GPX4) protects cells from ferroptosis by using glutathione (GSH) to reduce phospholipid hydroperoxides ^1, 2^, GPX4 production depends on the specialized machinery that incorporates selenium into proteins as selenocysteine (Sec). Unlike canonical amino acids, Sec is synthesized on a dedicated tRNA and inserted at UGA codons through translational recoding ^3–5^. Selenophosphate synthetase 2 (SEPHS2) supplies monoselenophosphate, the activated selenium donor required for Sec synthesis. The dependence of GPX4 on this pathway provides a rationale for targeting selenoprotein biosynthesis to induce ferroptosis. We therefore reasoned that disrupting SEPHS2 would compromise GPX4-mediated lipid protection and primarily induce ferroptotic cell death.

GPX4-dependent protection also requires a sufficient supply of cysteine for GSH synthesis. The cystine/glutamate antiporter system x_c_^⁻^, whose substrate transport is mediated by SLC7A11 (xCT), supports this requirement by importing extracellular cystine. However, imported cystine must first be reduced to cysteine, imposing a demand on cellular reducing capacity. Cystine uptake therefore supports antioxidant defense only when cells can accommodate its disulfide burden. This constraint is illustrated by disulfidptosis, a form of cell death under glucose limitation in which NADPH depletion in SLC7A11-high cells promotes aberrant disulfide bonding in cytoskeletal proteins, actin network collapse, and cell death ^6^. Thus, increasing the supply of a nutrient used for antioxidant defense can become detrimental when its reduction is limiting.

Here, we targeted SEPHS2 to impair GPX4 and induce ferroptosis. Unexpectedly, ferroptosis inhibition suppressed lipid peroxidation but only partially rescued cell viability. We found that SEPHS2 loss also compromised thioredoxin reductase 1 (TXNRD1)-dependent cystine reduction, which promoted NRF2-dependent SLC7A11 induction further increasing cystine influx. This imbalance generated disulfide stress and, upon GSH depletion, molecular features of disulfidptosis. In orthotopic neuroblastoma xenografts, SEPHS2 depletion increased disulfide-linked protein aggregation and suppressed tumor growth. Forced cystine uptake through SLC7A11 overexpression alone likewise suppressed xenograft growth, despite increased tumor GSH. These findings place selenoprotein biosynthesis at the crossroads of ferroptosis and disulfidptosis, revealing that a strategy intended to disable GPX4 also compromises the capacity to tolerate the cystine that sustains antioxidant defense.

## Results

### Disruption of selenoprotein biosynthesis compromises redox homeostasis beyond ferroptosis

Strategies to induce ferroptosis have attracted considerable interest as a means to selectively target specific tumor entities. Motivated by the high co-essentiality between GPX4 and the selenocysteine biosynthesis pathway (Fig. 1a, b), we sought to explore to what extend this pathway could be used to promote ferroptosis. Accordingly, genetic loss of members of the selenoprotein incorporation pathway (SEPHS2, EEFSEC, or SECISBP2) in HT1080 fibrosarcoma cells markedly reduced selenoprotein expression, including GPX1, GPX4, and SELENON, and cell viability (Fig. 1c and Extended Data Fig. 1a, b).

**Fig. 1.**
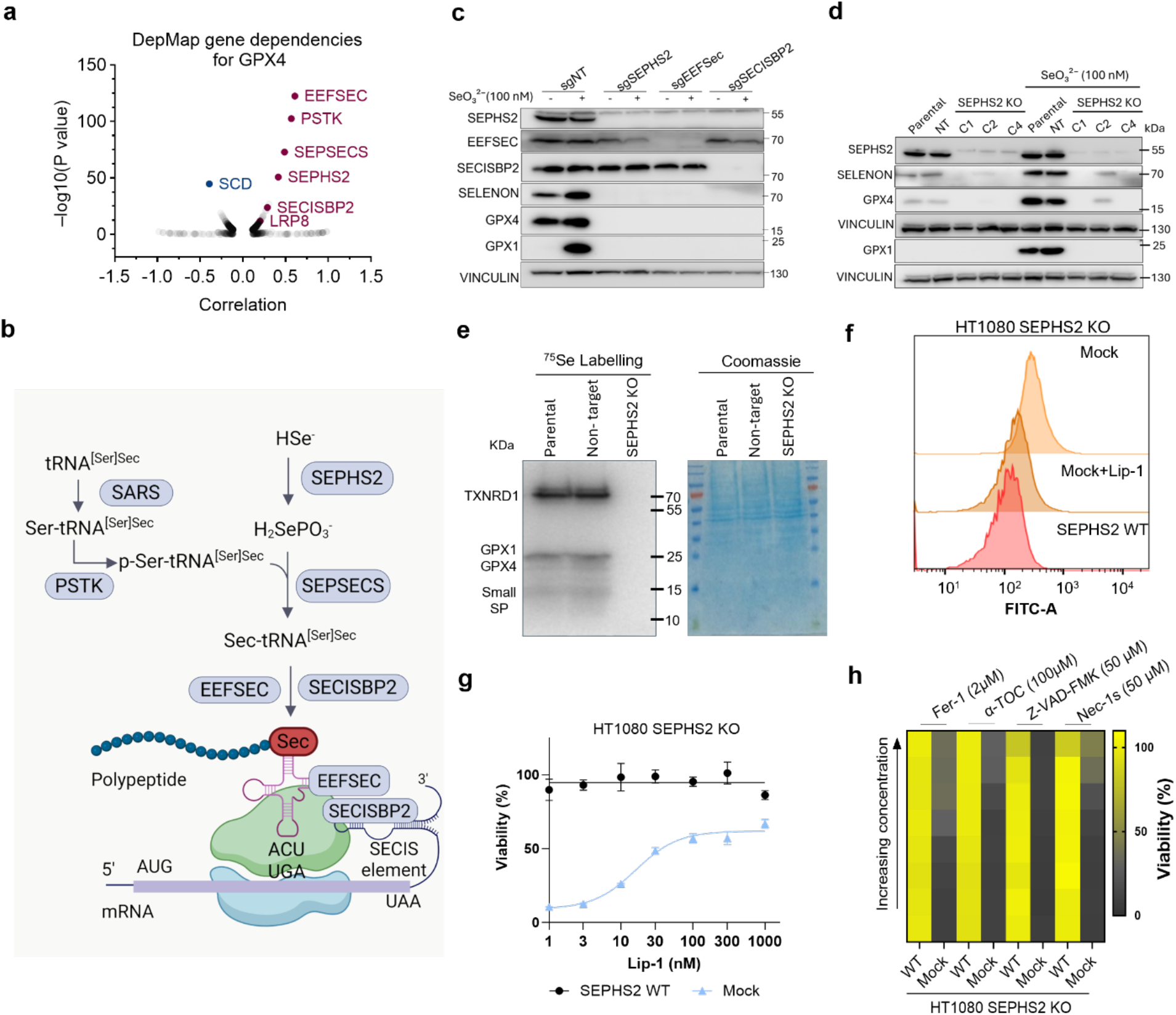
Disruption of selenoprotein biosynthesis compromises redox homeostasis beyond ferroptosis. a, Gene-dependency analysis for GPX4. Data are extracted from https://depmap.org. **b,** Schematic representation of the selenoprotein biosynthesis pathway. **c,** Immunoblot analysis of selenoproteins after knocking out SEPSECS, EEFSEC, and SEPHS2 in HT1080 cells with and without 100 nM selenite. **d,** Immunoblot analysis selenoproteins in SEPHS2-KO HT1080 single-cell clones (C1, C2 and C4), with and without 100nM selenite. **e,** ^75^Se radioisotope labelling to assess selenoprotein biosynthesis in SEPHS2-KO C1 cells. **f,** Measuring lipid peroxidation using C11-BODIPY 581/591 staining in WT or SEPHS2-KO cells, with or without treatment with 500 nM Lip-1. **g,** Viability of SEPHS2-KO HT1080 cells reconstituted with SEPHS2 WT or an empty vector (Mock), in the presence of the increasing concentrations of Lip-1. Data plotted are mean ± s.d. of triplicate wells. **h,** Viability of SEPHS2-WT or-KO HT1080 cells in the presence of increasing concentrations of the indicated cell death inhibitors. Highest used concentration is indicated. Data plotted are mean ± s.d. of triplicate wells.

We next focused on SEPHS2, which catalyzes the synthesis of monoselenophosphate from selenide and ATP and provides the only known selenium donor required for selenocysteine biosynthesis. We hypothesized that SEPHS2 loss would broadly compromise global selenoprotein production and thereby impair cellular resistance to ferroptosis. Therefore, we generated clonal SEPHS2-KO cells and confirmed that, unlike controls, they could not synthesize selenoproteins after selenium supplementation (Fig. 1d). Consistent with SEPHS2’s central role in selenium utilization, radiolabeled selenium tracing showed that SEPHS2-deficient cells failed to incorporate selenium into selenoproteins (Fig. 1e).

Because SEPHS2 itself contains a Sec, we next investigated the contribution of this residue to its function. As expected, re-expression of wild-type (WT) SEPHS2 rescued the viability of SEPHS2-deficient cells. Similarly, the U60C variant, in which Sec is replaced by cysteine, provided partial rescue of selenoprotein expression and cell viability (Extended Data Fig. 1c). As previously reported for other selenoproteins, expression of the cysteine variant accumulated to substantially higher levels than selenocysteine-containing SEPHS2, in line with the greater translational efficiency of cysteine relative to Sec^7^. Although the activity of cysteine mutant is expected to be lower than WT ^8^, higher abundance likely compensates for the loss of activity. By contrast, the U60A variant failed to restore selenoprotein expression or cell viability (Extended Data Fig. 1d-f). We next assessed the sensitivity of cells expressing WT or U60C variants of SEPHS2 to the ferroptosis inducers RSL3, erastin, and buthionine sulfoximine (BSO). Consistently, expression of both the WT and U60C variants conferred a similar degree of resistance despite the difference in protein abundance (Extended Data Fig. 2a, b).

Because SEPHS2 overexpression increased GPX4 abundance and enhanced resistance to RSL3, erastin and BSO (Extended Data Fig. 2c, d), we next determined whether SEPHS2 deficiency exclusively affected ferroptosis sensitivity. Given GPX4’s essential role in preventing ferroptosis, SEPHS2-KO cells required constant supplementation with the ferroptosis inhibitor Liproxstatin-1 (Lip-1) to maintain viability and enable propagation in culture. Consistent with a role in ferroptosis, SEPHS2-KO cells showed increased lipid peroxidation, which was completely abrogated by Lip-1 treatment (Fig. 1f and Extended Data Fig. 2e). However, treatment with Lip-1 could only partially restore cell viability and did not fully rescue cell fitness (Fig. 1g). A broader assessment of cell-death inhibitors similarly identified ferroptosis inhibition as the only intervention that partially rescued the phenotype under the conditions tested (Fig. 1h). Thus, SEPHS2 deficiency induces a ferroptotic component of cell death but also causes additional stress that is not explained by ferroptosis alone.

### SEPHS2 deficiency induces SLC7A11-dependent cystine toxicity

Given that previous reports suggested that loss of SEPHS2 could promote toxicity due to increase production of selenide derived from selenite ^8^, we investigated if that could be a potential mechanism. Nevertheless, we did not observe additional toxicity in the absence of SEPHS2, when we treated cells with increasing concentrations of selenite or selenocystine (Extended Data Fig. 2f), arguing against a relevant role of this enzyme in the detoxification of selenide under the conditions assayed.

To identify pathways that might account for the additional toxicity associated with SEPHS2 loss, we performed quantitative proteomic analysis of isogenic SEPHS2-deficient cells reconstituted with either WT SEPHS2 or an empty-vector control. SEPHS2 deficiency caused a clear difference in the proteome of the KO cells compared with WT cells and reduced the abundance of multiple selenoproteins (Fig. 2a and Extended Data Fig. 3a). The extent of this reduction varied among selenoproteins, with particularly pronounced changes among proteins whose Sec residues occur closer to the protein N-terminal sequence (Extended Data Fig. 3b). Pathway enrichment analysis confirmed that upregulated proteins were enriched in NRF2-dependent antioxidant signaling, ferroptosis-related, and nuclear receptor pathways, while downregulated proteins were significantly enriched in mitochondrial oxidative phosphorylation and electron transport chain pathways, indicative of mitochondrial disfunction (Extended Data Fig. 3c).

**Fig. 2.**
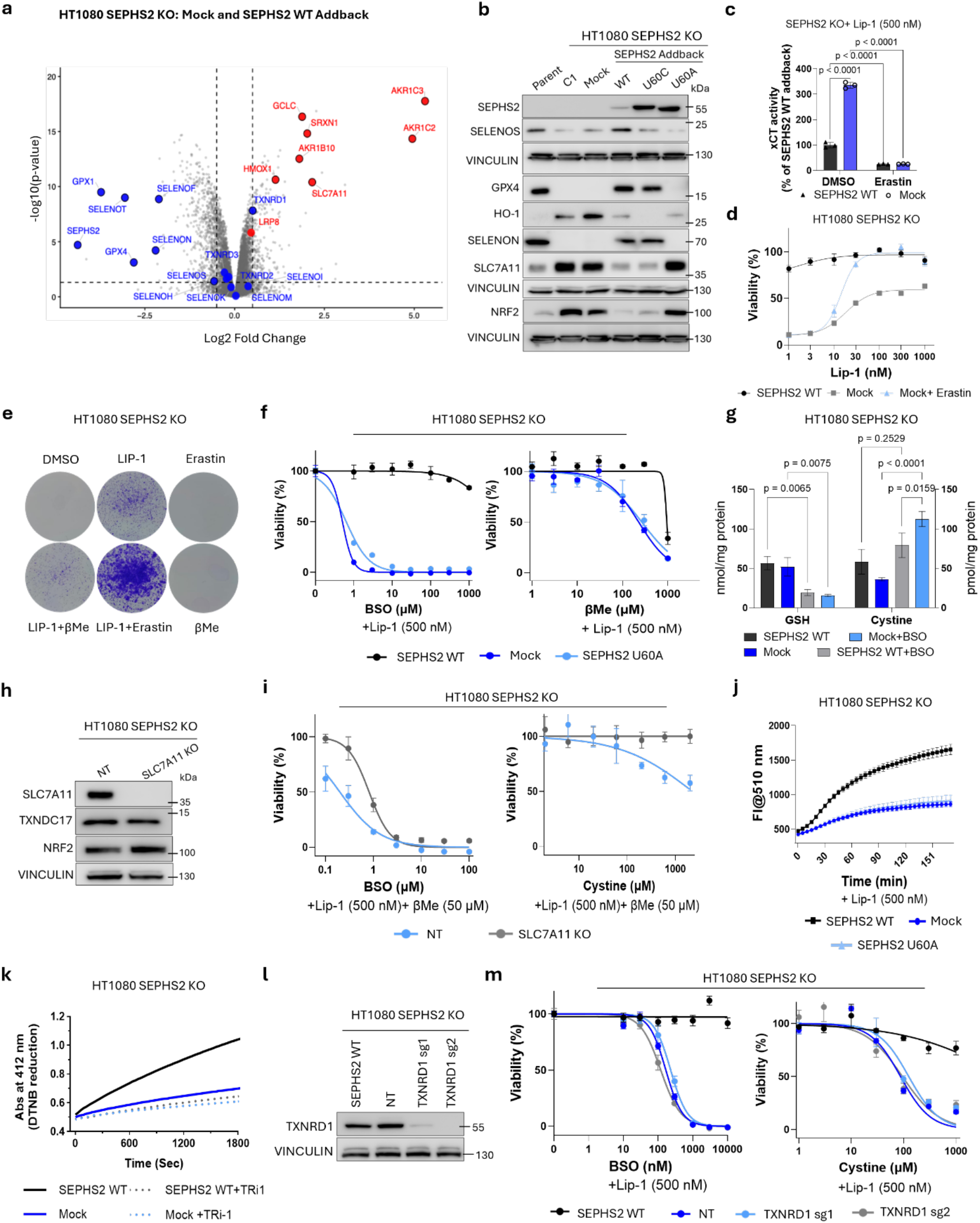
SEPHS2 deficiency promotes cystine-associated stress and alters cellular disulfide-reducing capacity. a, Proteomic analysis of SEPHS2-KO HT1080 cells reconstituted with SEPHS2 WT or the mock control. Detected selenoproteins are shown in blue. NRF2 target genes are shown in red. **b,** Immunoblot analysis of SEPHS2-KO cells reconstituted with SEPHS2 WT or the indicated SEPHS2 mutants. **c,** Analysis of SLC7A11 transport activity in SEPHS2-KO HT1080 cells reconstituted with SEPHS2 WT or the mock control. Data plotted are mean ± s.d. of triplicate wells. **d,** Viability of SEPHS2-KO HT1080 cells reconstituted with SEPHS2 WT or the mock control, following treatment with 250 nM erastin and the increasing concentrations of Lip-1. Data plotted are mean ± s.d. of triplicate wells. **e,** Clonogenecity of SEPHS2-KO cells upon treatment with the indicated supplementation indicated. **f,** Viability of SEPHS2-KO HT1080 cells reconstituted with SEPHS2 WT, the mock control, or the inactive U60A mutant construct in response to increasing concentrations of BSO (left) or βME (right). Data plotted are mean ± s.d. of triplicate wells. **g,** Quantification of intracellular reduced GSH and cystine levels in SEPHS2-KO HT1080 cells reconstituted with SEPHS2 WT or the mock control, with or without 200 μM BSO treatment, using targeted LC-MS/MS. Data plotted are mean ± s.d. of triplicate wells. **h,** Immunoblot analysis of SEPHS2-KO cells transduced with a non-target or SLC7A11-targetting sgRNA. **i,** Viability of SEPHS2-KO HT1080 cells transduced with a non-target or SLC7A11-targetting sgRNA in response to increasing concentrations of BSO (left) or cystine (right). Data plotted are mean ± s.d. of triplicate wells. **j,** Fluorescence intensity of BODIPY-cystine in SEPHS2-KO cells reconstituted with SEPHS2 WT, the mock control, or the inactive U60A mutant construct. Data plotted are mean ± s.d. of triplicate wells. **k,** DTNB reduction assay to determine the disulfide-reducing capacity of cell lysates from SEPHS2-KO HT1080 cells reconstituted with SEPHS2 WT or the mock control, with or without treatment with 1μM of the TXNRD1 inhibitor, TRi-1. Data plotted are mean of technical triplicate well. **l,** Immunoblot analysis of SEPHS2-KO cells transduced with a non-target or TXNRD1-targetting sgRNA. **m**, Viability of SEPHS2-KO HT1080 cells transduced with a non-target or TXNRD1-targetting sgRNA in response to increasing concentrations of BSO (left) or cystine (right). Data plotted are mean ± s.d. of triplicate wells.

We first confirmed that SEPHS2 deficiency was associated with increased abundance of proteins linked to NRF2 signaling by immunoblotting (Fig. 2a, b). This effect was not restricted to SEPHS2, as we observed NRF2 upregulation after disrupting other components of the selenoprotein biosynthesis machinery, including EEFSEC and SECISBP2, reflecting a broader consequence of impaired selenoprotein biosynthesis (Extended Data Fig. 3d).

Among the upregulated proteins, we focused on SLC7A11, the substrate-transporting subunit of system x_c_^−^, which mediates cellular cystine uptake in exchange for glutamate ^9, 10^ and is known to protect cells from ferroptosis and oxidative stress. SLC7A11 abundance and transport activity were both increased in SEPHS2-deficient cells (Fig. 2b, c). We hypothesized that enhanced cystine uptake represented a compensatory response to SEPHS2 loss and that inhibiting this response would exacerbate the viability defect. Unexpectedly, inhibition of system x_c_^−^ with a subtoxic concentration of erastin in the presence of Lip-1 improved cell viability more effectively than Lip-1 alone (Fig. 2d, e and Extended Data Fig. 3e). Combined Lip-1 and erastin treatment also enhanced cell survival and proliferation, whereas supplementation with β-mercaptoethanol (βME) in the presence of Lip-1 impaired proliferation (Extended Data Fig. 3f). These observations prompted us to consider whether increased cystine uptake instead imposed a burden on the compromised cellular disulfide-reducing capacity. Consistent with this possibility, treatment with tris(2-carboxyethyl)phosphine (TCEP), a reducing agent that promotes extracellular cystine reduction, rescued SEPHS2-deficient cells to an extent comparable to erastin treatment (Extended Data Fig. 3g, h). Together, these findings suggest that increased cystine uptake contributes to the Lip-1-insensitive component of the SEPHS2-deficient phenotype, implicating impaired cystine metabolism and thiol–disulfide homeostasis as additional sources of cellular stress.

Previous studies have linked inhibition of the selenoprotein TXNRD1 to cell death; therefore, loss of SEPHS2 could also impair thioredoxin-dependent redox homeostasis by compromising the insertion of selenocysteine in one of the redox centers of this enzyme ^11^. Motivated by this, we next tested whether SEPHS2-deficient cells were vulnerable to perturbations that increase cystine-associated stress or limit cellular reducing capacity. Proteomic analysis also revealed increased abundance of glutamate–cysteine ligase catalytic subunit (GCLC), which catalyzes the rate-limiting step in GSH synthesis (Fig. 2a). Given the potential impairment of TXNRD1-dependent redox regulation following SEPHS2 loss, we hypothesized that the GSH-dependent reducing system might serve as an alternative mechanism for maintaining cellular thiol-disulfide homeostasis. We therefore examined the effects of BSO, a GCLC inhibitor, and βME, which alters extracellular cystine availability and cellular cysteine supply. Cells reconstituted with WT SEPHS2 were more resistant to both treatments than cells expressing the empty-vector control or the inactive U60A variant (Fig. 2f and Extended Data Fig. 3i). Targeted sulfur metabolome analysis revealed that BSO treatment markedly depleted intracellular GSH and increased cystine accumulation in both SEPHS2-sufficient and-deficient cells. Notably, cystine accumulation was more pronounced in SEPHS2-deficient cells, potentially contributing to their increased susceptibility to cystine-associated toxicity (Fig. 2g and Extended Data Fig. 4a).

Erastin also increased resistance to BSO, further linking the observed vulnerability to system x_c_^−^-dependent cystine uptake (Extended Data Fig. 4b). To test this relationship genetically, we deleted SLC7A11 in SEPHS2-deficient cells (Fig. 2h). SLC7A11 loss increased resistance to exogenous cystine, BSO and βMe (Fig. 2i, and Extended Data Fig. 4c). Extracellular pyruvate supplementation also conferred mild resistance to BSO-associated toxicity (Extended Data Fig 4d), potentially by sequestering excess cysteine through the formation of reversible cysteine–pyruvate conjugates, as recently described in cancer cells ^12^. Together, these findings establish that increased SLC7A11-dependent cystine uptake contributes to the sensitivity of SEPHS2-deficient cells to cystine-associated stress and GSH depletion. These data also establish that GSH-dependent reducing mechanisms partially compensates for impaired TXNRD1 function.

### SEPHS2 loss impairs TXNRD1, contributing to reduced cellular disulfide-reducing capacity

We next investigated why increased cystine uptake was detrimental in SEPHS2-deficient cells. Because TXNRD1 is a selenoprotein required to maintain the thioredoxin system and cellular redox homeostasis ^11, 13, 14^, we examined whether impaired TXNRD1 function could contribute to this phenotype.

Using a kinetic assay based on fluorescently labeled cystine, we found that SEPHS2-deficient cells reconstituted with WT SEPHS2 reduced the substrate, whereas cells expressing the empty vector or the U60A variant showed substantially impaired cystine-reducing activity (Fig. 2j). We then assessed the disulfide-reducing capacity of cell lysates using the DTNB reduction assay. In cells lacking functional SEPHS2, treatment with the TXNRD1 inhibitor TRi-1 did not further reduce the measured activity, whereas TRi-1 decreased reducing capacity in cells reconstituted with WT SEPHS2 (Fig. 2k). These findings are consistent with the specific impairment of TXNRD1-dependent reducing capacity following SEPHS2 loss.

To investigate whether the truncated TXNRD1 protein retained any residual function in maintaining redox homeostasis following SEPHS2 loss, we generated TXNRD1 KO cells in a SEPHS2-deficient background (Fig. 2l). TXNRD1 deletion did not further increase sensitivity to BSO, cystine or βMe treatment (Fig. 2m, Extended Data Fig. 4E). These findings demonstrate that impaired Sec incorporation following SEPHS2 deficiency is sufficient to disrupt TXNRD1-dependent redox function. They are also consistent with SEPHS2 deficiency and TXNRD1 loss affecting a shared pathway that limits cellular tolerance to cystine-associated stress.

We next tested whether TXNDC17 overexpression could rescue cystine-associated toxicity in SEPHS2-deficient cells. TXNDC17 is a cytosolic oxidoreductase that catalyzes cystine reduction to cysteine using TXNRD1-and NADPH-derived reducing equivalents ^15^. TXNDC17 overexpression failed to restore cystine tolerance (Extended Data Fig. 4f, g), suggesting that increasing TXNDC17 abundance is insufficient when TXNRD1-dependent reducing capacity is compromised. These data confirm the dependence of TXNDC17-mediated cystine reduction on functional TXNRD1.

Collectively, these findings indicate that SEPHS2 deficiency compromises two complementary redox-protective systems: GPX4-dependent protection against lipid peroxidation and TXNRD1-associated disulfide-reducing capacity. In parallel, increased SLC7A11-dependent cystine uptake, which usually protects against ferroptosis, becomes detrimental when disulfide-reducing capacity is impaired and exacerbates the consequences of disrupted redox homeostasis. This combination provides a mechanistic explanation for why ferroptosis inhibition alone does not fully rescue the loss of viability caused by SEPHS2 deficiency.

### NRF2-dependent SLC7A11 induction amplifies cystine-associated stress

We next investigated the mechanism responsible for SLC7A11 upregulation in SEPHS2-deficient cells. Because NRF2 signature was elevated in the proteomic analysis, we generated SEPHS2– NRF2 double-knockout cells and reconstituted them with SEPHS2, NRF2, both proteins, or the corresponding empty constructs (Fig. 3a). These experiments showed that SLC7A11 upregulation in SEPHS2-deficient cells was dependent on NRF2. Consistent with a detrimental contribution of NRF2-dependent SLC7A11 induction under these conditions, NRF2 loss increased the resistance of SEPHS2-deficient cells to BSO or cystine treatment (Fig. 3b).

**Fig. 3.**
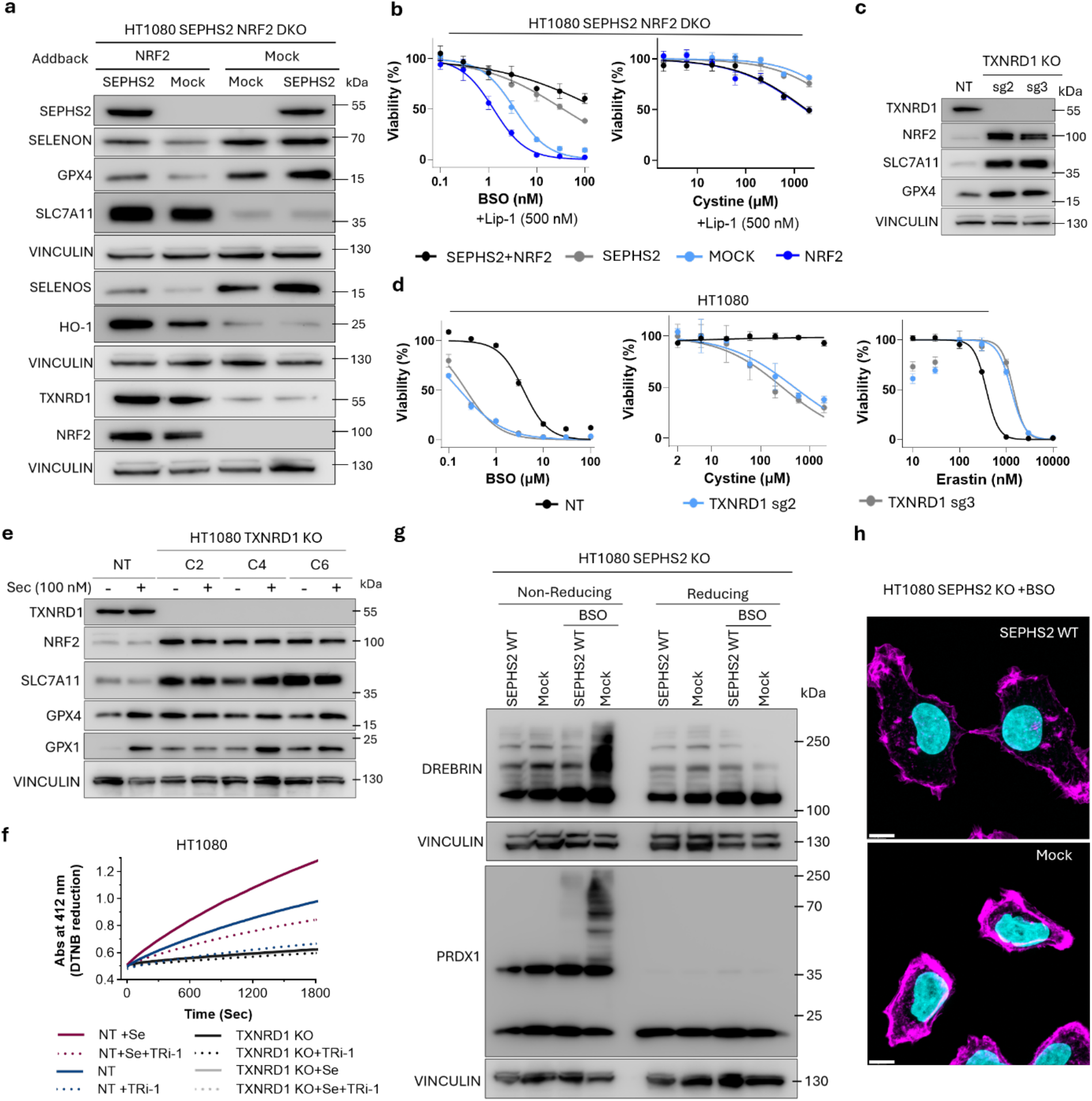
NRF2-dependent SLC7A11 induction amplifies selenoprotein loss induced disulfidptosis. a, Immunoblot analysis of SEPHS2/NRF2 DKO cells reconstituted with SEPHS2, NRF2, both proteins, or respective controls. **b,** Viability of SEPHS2/NRF2 DKO cells reconstituted with SEPHS2, NRF2, both proteins, or respective controls in response to increasing concentrations of BSO (left) or cystine (right). Data plotted are mean ± s.d. of triplicate wells. **c,** Immunoblot analysis of TXNRD1-KO HT1080 cells. **d,** Viability of HT1080 cells transduced with non-target or two sgRNAs targeting TXNRD1 in response to increasing concentrations of BSO (left), cystine (middle), or erastin (right). Data plotted are mean ± s.d. of triplicate wells. **e,** Immunoblot analysis of single clone TXNRD1-KO HT1080 cells, with or without 100 nM selenocystine. **f,** DTNB reduction assay to determine the disulfide-reducing capacity of cell lysates from parental or TXNRD1-KO HT1080 cells, with or without treatment with 100 nM selenite and 1 µM of the TXNRD1 inhibitor, Tri-1. Data plotted are mean of duplicates. **g,** Immunoblot analysis to detect aggregates of Drebrin and PRDX1 under non-reducing or reducing conditions in SEPHS2-knockout cells reconstituted with SEPHS2 WT or mock control. **h,** Phalloidin staining (magenta) to reveal F-ACTIN aggregates in SEPHS2-knockout cells reconstituted with SEPHS2 WT or mock control. DNA is counterstained with DAPI (cyan). Scale bar is 10 µm.

Interestingly, SEPHS2 deficiency was associated with increased TXNRD1 expression in NRF2-proficient cells (Fig. 3a), despite the evidence for impaired disulfide-reducing capacity. NRF2 activation likely drives increased TXNRD1 expression; however, impaired Sec biosynthesis prevents the incorporation of Sec into TXNRD1, resulting in protein that lacks the essential catalytic residue required for disulfide-reducing activity. Thus, increased TXNRD1 protein abundance does not translate into increased enzymatic activity.

Consistent with a link between TXNRD1 activity loss and NRF2 activation ^16^, genetic deletion of TXNRD1 increased NRF2 and consequently SLC7A11 abundance (Fig. 3c). TXNRD1-deficient cells were also more sensitive to BSO or cystine treatment while showing increased resistance to erastin and RSL3 (Fig. 3d and Extended Data Fig. 4h). These findings support a model in which impaired TXNRD1-dependent reducing capacity activates an NRF2-associated response that induces increased SLC7A11 expression. Although this response may be adaptive in some contexts, increased cystine uptake becomes detrimental when the capacity to maintain intracellular thiol–disulfide balance is compromised.

We next investigated whether increasing the expression of other selenoproteins in TXNRD1-deficient cells could attenuate NRF2 activation. Although Sec supplementation increased the expression of other selenoproteins, it failed to reduce NRF2 activation (Fig. 3e), suggesting that TXNRD1 loss is a major driver of this response and cannot be compensated for by restoring the expression of other selenoproteins under the conditions tested. Consistent with this, TXNRD1-deficient cells showed markedly reduced DTNB-reducing activity, which Sec supplementation did not restore. In contrast, Sec supplementation enhanced DTNB reduction in WT cells, whereas treatment with the TXNRD1 inhibitor TRi-1 suppressed this activity (Fig. 3f). These findings identify TXNRD1 as a major contributor to cellular disulfide-reducing capacity and suggest that its loss compromises protection against disulfide stress.

Motivated by these observations, we explored whether increased cystine uptake and impaired disulfide-reducing capacity were associated with molecular features of disulfidptosis in SEPHS2-deficient cells. Following BSO treatment, we assessed Drebrin oligomerization, PRDX1 dimerization and F-actin reorganization (Fig. 3g, h). Non-reducing immunoblotting revealed increased high-molecular-weight Drebrin and PRDX1 species, which disappeared upon βME treatment, consistent with disulfide-linked protein complexes (Fig. 3g). In parallel, BSO-induced GSH depletion caused F-actin contraction and cytoskeletal reorganization, recapitulating features of disulfidptosis (Fig. 3h and Extended Data Fig. 4i). Together, these findings demonstrate that GSH depletion promotes disulfide-dependent protein aggregation and actin cytoskeletal abnormalities in SEPHS2-deficient cells, although the contribution of these changes to cystine-associated toxicity remains to be established.

### SEPHS2 deficiency extends cell death vulnerability to neuroblastoma

Next, we asked whether the effects of SEPHS2 deficiency observed in HT1080 cells were also evident in neuroblastoma models, a cancer context in which dependence on selenium utilization and GPX4 has previously been established ^17, 18^. In SK-N-DZ neuroblastoma cells, SEPHS2 deficiency reduced selenoprotein abundance and increased NRF2 and its target proteins (Fig. 4a). To establish whether these changes were reversible, we generated SEPHS2-deficient SK-N-DZ cells carrying a doxycycline (DOX)-inducible WT SEPHS2 construct. Induction of SEPHS2 expression restored the altered protein-expression profile, whereas Dox withdrawal caused the cells to revert towards the SEPHS2-deficient state over the following days including loss of selenoproteins and NRF2 activation (Fig. 4b and Extended Data Fig. 5a, b). These findings confirm the link between the observed molecular changes and SEPHS2 expression. Upon DOX withdrawal, the cells were also more sensitive to BSO treatment (Fig. 4c), extending the association between impaired selenoprotein biosynthesis and cystine-associated vulnerability to a second cancer model.

**Fig. 4.**
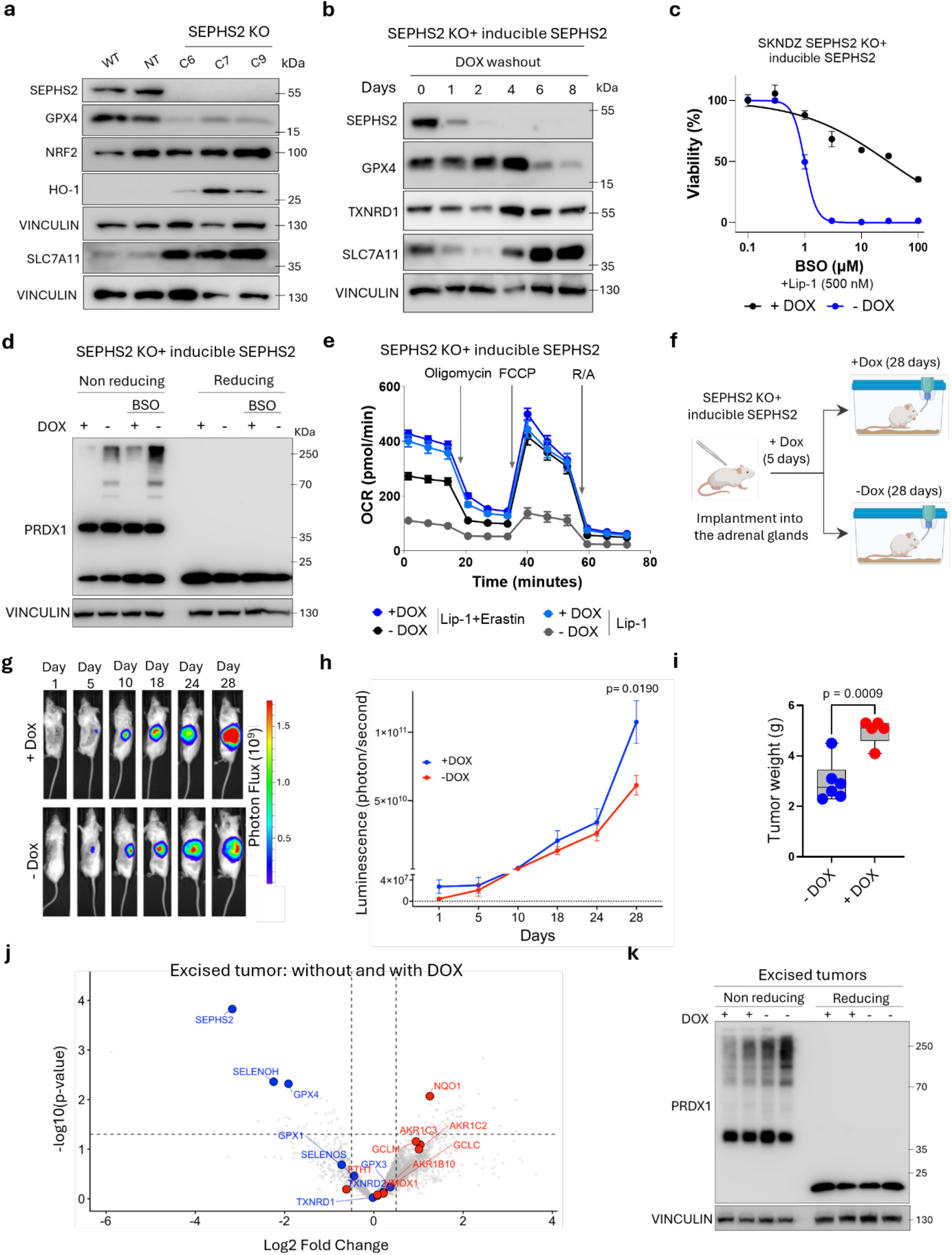
SEPHS2 sustains redox homeostasis and tumor growth in neuroblastoma. a, Immunoblot analysis of independent SEPHS2-KO SK-N-DZ single-cell clones (C6, C7 and C9).**b,** Immunoblot analysis of SEPHS2-KO SK-N-DZ cells, reconstituted with DOX-inducible SEPHS2. Following induction with 500 ng/ml, DOX was withdrawn to assess SEPHS2 protein depletion and the accompanying recovery of SLC7A11 expression. **c,** Viability of SEPHS2-KO SK-N-DZ cells, reconstituted with DOX-inducible SEPHS2 with or without DOX treatment, in the presence of increasing concentrations of BSO. Data plotted are mean ± s.d. of triplicate wells. **d,** Immunoblot analysis to detect aggregates of Drebrin under non-reducing or reducing conditions in SEPHS2-KO SK-N-DZ cells, reconstituted with DOX-inducible SEPHS2 with or without treatment with 500 ng/ml of DOX. **e,** Mitochondrial OCR in SEPHS2-KO SK-N-DZ cells, reconstituted with DOX-inducible SEPHS2 or mock control, with or without treatment with 500 ng/ml of DOX and 500 nM Lip-1, in the presence or absence of 250 nM of erastin. Data plotted are mean ± s.d. of triplicate wells. **f,** Experimental design of the injection of luciferase expressing SEPHS2-KO SK-N-DZ cells, reconstituted with DOX-inducible SEPHS2 into the adrenal glands of mice. **g,** Representative luminescence images of mice in F on the indicated days post injection. **h,** Quantification of the luminescence signal as an indication of tumor volume. Data plotted are mean ± s.d. of n= 6 mice per group. **i**, Quantification of tumor volume from the tumors extracted from the mice in **a** after scarification on day 28. Data plotted are mean ± s.d. of at least n= 5 mice. **j,** Proteomics analysis of excised tumor treated without or with 2mg/mL DOX treatment in the drinking water. Detected selenoproteins are shown in blue. NRF2 target genes are shown in red. **k,** Immunoblot analysis of PRDX1 under non-reducing and reducing conditions in excised tumor lysates with or without DOX (2 mg/mL) treatments

We next examined molecular features associated with disulfidptosis in SEPHS2-deficient SK-N-DZ cells. BSO treatment induced disulfide-linked high-molecular-weight species of PRDX1 in SEPHS2-deficient cells following Dox withdrawal, an effect that disappeared upon βME treatment (Fig. 4d).

Because an excess of sulfur containing amino acids has been linked to mitochondrial dysfunction and impaired respiration ^19–21^, we next measured oxygen consumption rates (OCR) in SK-N-DZ cells under SEPHS2-expressing (in the presence of DOX) and SEPHS2-deficient (in the absence of DOX) conditions. SEPHS2 deficiency reduced mitochondrial respiration, which was rescued by erastin treatment (Fig. 4e and Extended Data Fig. 5c). These data reveal that in SEPHS2-deficient cells excessive SLC7A11-mediated cystine uptake contributes to impaired mitochondrial respiration.

To explore the consequences of altered SEPHS2 expression *in vivo*, we used a previously reported orthotopic neuroblastoma model ^17^. SEPHS2-deficient SK-N-DZ cells carrying inducible WT SEPHS2 were implanted into the adrenal gland of mice, and SEPHS2 expression was regulated through DOX administration. On day 5, the mice were randomly assigned to two groups with either continuation or stopping DOX treatment (Fig. 4f). Withdrawal of DOX resulted in a significant delay in tumor growth, as reflected by reduced bioluminescence signals during tumor progression and lower tumor weights at sacrifice on day 28 post-injection (Fig. 4g-i and Extended Data Fig. 5d). Proteomics and immunoblot analysis of excised tumor tissues confirmed reduced selenoprotein expression following DOX withdrawal (Fig. 4j and Extended Data Fig. 5e). Furthermore, non-reducing immunoblotting revealed increased high-molecular-weight PRDX1 species, consistent with enhanced disulfide-linked protein aggregates (Fig. 4k). Together, these findings indicate that impaired selenoprotein biosynthesis compromises neuroblastoma tumor growth in vivo and is accompanied by molecular evidence of increased disulfide stress.

### SLC7A11 overexpression creates a vulnerability partially mitigated by TXNDC17

*MYCN*-amplified SK-N-DZ neuroblastoma cells generally exhibit low SLC7A11 expression (Fig. 4a), which may reflect a selective pressure against increased cystine uptake in a lineage particularly vulnerable to disulfide stress. We therefore asked whether restoring SLC7A11 expression would compromise neuroblastoma cell fitness and whether increased disulfide-reducing capacity could counteract its effects. Because erastin restored mitochondrial respiration in SEPHS2-deficient cells, we also investigated whether increased cystine uptake contributes to mitochondrial dysfunction. To this end, we overexpressed SLC7A11 in SK-N-DZ neuroblastoma cells, alone or together with TXNDC17 (Fig. 5a). SLC7A11 overexpression increased sensitivity to BSO and βME, whereas co-expression of TXNDC17 restored resistance (Fig. 5b). Supplementation with Sec only partly rescued this sensitivity, suggesting that TXNRD1-dependent reducing capacity contributes to tolerance of cystine-associated stress. Consistent with this, under partial GSH depletion with a low concentration of BSO (1 µM), DOX-induced SLC7A11-overexpressing and mock cells showed reduced proliferation, whereas co-expression of TXNDC17 restored proliferation (Fig 5c, d). These findings suggest that increased TXNDC17 expression can support cell fitness when GSH availability is limited.

**Fig. 5.**
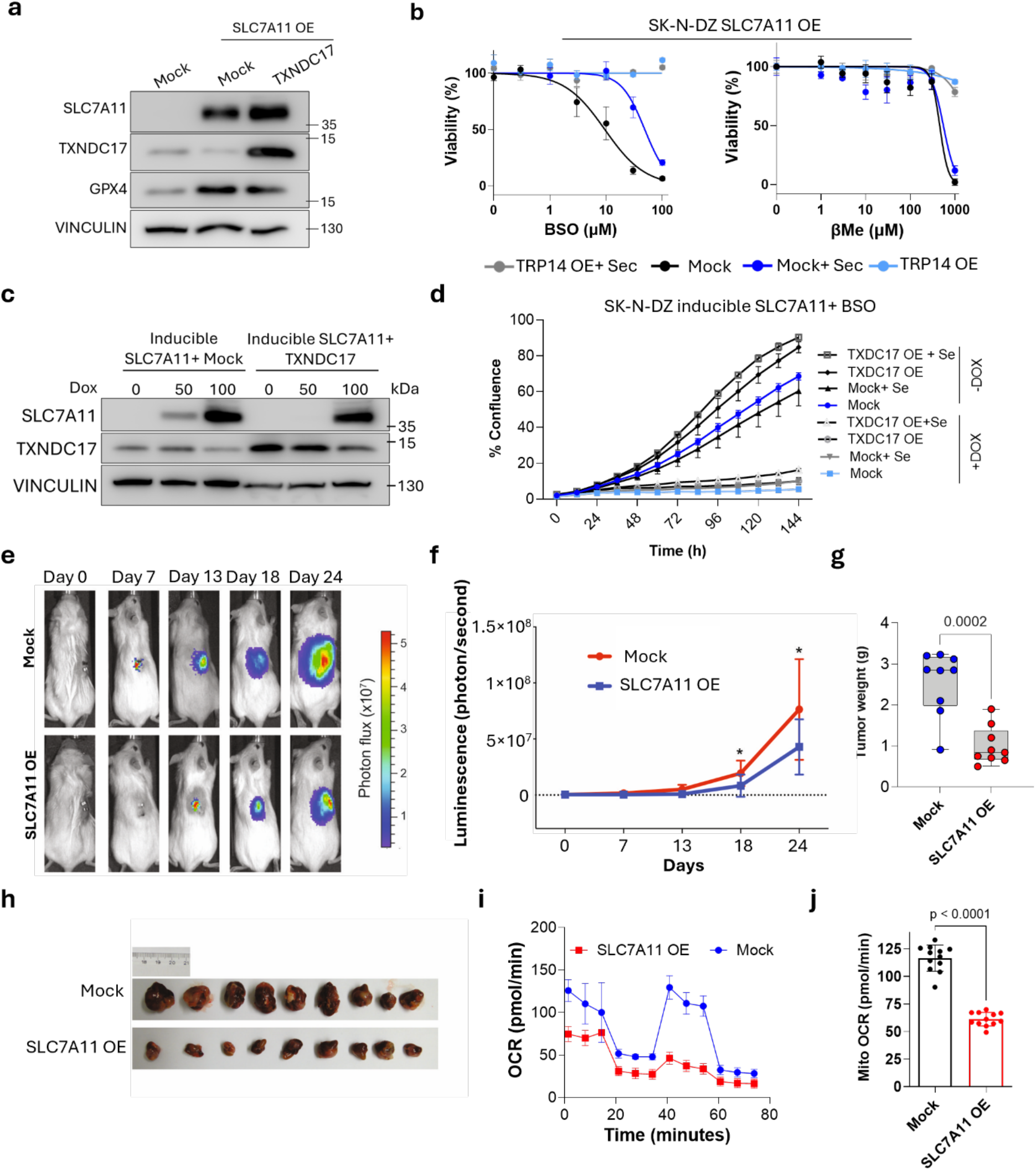
SLC7A11 overexpression damages the cellular fitness of SKNDZ neuroblastoma. a, Immunoblot analysis of SK-N-DZ cells transduced with an empty mock or SLC7A11 overexpressing construct, with or without co-expression of TXNDC17. **b,** Viability of SK-N-DZ cells transduced with an empty mock or SLC7A11 overexpressing construct, with or without co-expression of TXNDC17 upon treatment with increasing concentrations of BSO (left) of βME (right). Data plotted are mean ± s.d. of triplicate wells. **c**, Immunoblot analysis of SK-N-DZ cells transduced with DOX inducible SLC7A11 overexpressing construct, with or without co-expression of TXNDC17. **d,** Proliferation analysis of SK-N-DZ cells, transduced with an inducible SLC7A11 overexpressing construct treated with 500 ng/mLof DOX, with or without co-overexpression of TXNDC17, treated with 100 nM selenite. Data plotted are mean ± s.d. of triplicate wells. **e,** Representative bioluminescence images of mice bearing SK-N-DZ xenografts expressing mock or SLC7A11 overexpression construct at the indicated days post injection. **f,** Quantification of the luminescence signal as an indication of tumor volume Data plotted are mean ± s.d. of n= 9 mice per group. **g,** Quantification of tumor weight from the tumors extracted from the mice in A, after scarification on day 24. Data plotted are mean ± s.d. of n= 9 tumors per group. **h,** Representative images of excised tumors from mock and SLC7A11 overexpressing groups. **i,** Mitochondrial OCR in SK-N-DZ cells transduced with an empty mock or SLC7A11 overexpressing construct. Data plotted are mean ± s.d. of triplicate wells. **j**, Parameters of oxygen consumption rate analysis of mock and SLC7A11 overexpressing cells showing mitochondrial maximal respiration. Data plotted are mean ± s.d. of n= 10 repeats.

We next assessed the consequences of SLC7A11 overexpression for tumor growth in vivo. Orthotopic implantation of SLC7A11-overexpressing SK-N-DZ cells resulted in significantly reduced tumor growth from day 18 after implantation, as confirmed by endpoint tumor-volume measurements (Fig. 5e–h). Metabolomic profiling of these tumors revealed increased GSH and N-methyl-L-glutamate, together with increased cystathionine, suggesting changes in glutathione and sulfur amino acid metabolism (Extended Data Fig. 6a–e).

To investigate the metabolic basis of this phenotype, we measured oxygen consumption and performed stable-isotope tracing in SLC7A11-overexpressing cells. These cells exhibited reduced oxygen consumption (Fig. 5i, j and Extended Data Fig. 6f–h), consistent with previous reports linking cysteine overload to mitochondrial dysfunction and reductive stress ^19–21^. We next used ^13^C-glutamine tracing to examine changes in mitochondrial and biosynthetic metabolism (Extended Data Fig. 7a). Although intracellular glutamine labeling was largely preserved, the fractional abundance of m+5 glutamate and m+5 GSH increased, whereas total proline and its m+5 and m+3 isotopologues decreased (Extended Data Fig. 7b, c). Intracellular cystine increased, while cystathionine was substantially depleted, potentially reflecting increased cystathionine export through SLC7A11 and/or altered transsulfuration pathway activity (Extended Data Fig. 7d, e). Glutamine-derived labeling of TCA-cycle intermediates was also altered, with reduced m+5 α-ketoglutarate and decreased m+3 aspartate and malate, despite unchanged total aspartate abundance (Extended Data Fig. 7f–i). These changes indicate altered distribution of glutamine-derived carbon across mitochondrial and biosynthetic metabolite pools.

We then performed ^13^C-glucose tracing to assess glucose-derived carbon utilization (Extended Data Fig. 7j). Labeling of glucose and the glycolytic metabolites pyruvate, lactate and alanine was largely preserved (Extended Data Fig. 7k–m). In contrast, glucose-derived labelling of serine (m+3, m+2 and m+1) and glycine (m+2) was reduced (Extended Data Fig. 7n, o), potentially contributing to the reduced nucleotide triphosphate pools observed following SLC7A11 overexpression. Labelling of m+2 α-ketoglutarate and glutamate was also decreased, whereas malate and aspartate labelling was largely maintained, with only modest reductions in their m+3 isotopologues. Succinate isotopologue distribution was strongly altered (Extended Data Fig. 7p– t). Together, these tracing experiments reveal substantial changes in the distribution of glutamine-and glucose-derived carbon across mitochondrial and biosynthetic pathways, consistent with the reduced respiratory capacity of SLC7A11-overexpressing cells.

Together, these findings show that increased SLC7A11-mediated cystine uptake is accompanied by extensive metabolic remodeling that affects the distribution of glutamine-and glucose-derived carbon across mitochondrial and biosynthetic pathways. These changes, together with impaired mitochondrial respiration, suggest that excessive cystine uptake imposes a broader metabolic burden on cells, further linking SEPHS2-dependent selenoprotein biosynthesis to maintaining disulfide and metabolic homeostasis.

## Discussion

Our findings identify SEPHS2 deficiency as a source of cellular redox stress that extends beyond impaired GPX4-mediated protection against lipid peroxidation. As an essential selenoprotein, GPX4 uses GSH to reduce phospholipid hydroperoxides and prevent ferroptosis ^1, 2^. Consistent with this function, SEPHS2 deficiency impaired selenoprotein synthesis and increased ferroptotic sensitivity. However, although Lip-1 completely prevented lipid peroxidation, it only partially restored cell fitness, indicating that additional mechanisms contribute to the detrimental effects of SEPHS2 loss.

We identified cystine-associated disulfide stress as the contributing mechanism. SLC7A11-mediated cystine uptake normally supports GSH synthesis, but cystine must first be reduced to cysteine, placing a demand on cellular disulfide-reducing capacity. In SEPHS2-deficient cells, impaired selenoprotein synthesis compromises this capacity, while NRF2-dependent SLC7A11 induction increases cystine uptake. This combination may create a feed-forward process in which increased cystine influx further exacerbates disulfide stress. Thus, SLC7A11-mediated cystine uptake, although generally protective, can become detrimental when cellular reducing capacity is insufficient.

The thioredoxin system appears to contribute to this vulnerability, where the Sec-containing TXNRD1 sustains thioredoxin-dependent reducing capacity ^22^, while the oxidoreductase TXNDC17 promotes cystine reduction ^15, 23^. In cells with functional TXNRD1, increased TXNDC17 expression supported proliferation when GSH synthesis was partially restricted. However, TXNDC17 overexpression did not rescue cystine-associated toxicity following SEPHS2 loss, suggesting that increasing TXNDC17 abundance is insufficient when TXNRD1 is compromised.

SEPHS2-deficient cells also displayed molecular features associated with disulfidptosis, including disulfide-linked DREBRIN and PRDX1 species and altered F-actin organization. Although these features resemble those described in SLC7A11-high cells under glucose limitation ^6, 24^, the initiating stress differs: SEPHS2 deficiency impairs selenoprotein-dependent redox capacity while inducing NRF2–SLC7A11 signaling. We therefore interpret these findings as evidence of increased susceptibility to cystine-driven disulfide stress, accompanied by molecular features of disulfidptosis, rather than canonical disulfidptosis.

Cystine-associated stress was also linked to mitochondrial dysfunction and metabolic remodeling from oxidative phosphorylation to gluconeogenesis ^25^. SEPHS2 deficiency reduced mitochondrial respiration in neuroblastoma cells, which was restored by blocking cystine uptake. Similarly, SLC7A11 overexpression reduced oxygen consumption and altered glutamine-and glucose-derived carbon distribution across TCA-cycle intermediates, as well as serine and glycine. These observations suggest that increased cystine uptake is associated with broader metabolic changes, although the contribution of individual pathways to cellular toxicity remains to be established.

The relevance of SEPHS2-dependent selenoprotein synthesis to tumor fitness was supported by our neuroblastoma models, in which SEPHS2 depletion impaired tumor growth in vivo. This is consistent with the reported dependence of *MYCN*-amplified neuroblastoma on selenium uptake ^17^ and the requirement for SEPHS2 in acute myeloid leukemia ^26^. Our findings further suggest that tumor dependence on selenium utilization may reflect the need to maintain multiple redox defense systems. This may be particularly relevant in KEAP1-mutant lung adenocarcinomas, where constitutive NRF2 activation increases SLC7A11 expression ^27^. High SLC7A11 activity can also constrain mitochondrial metabolism by promoting glutamate export and limiting its availability for TCA-cycle anaplerosis ^28^, potentially adding to the metabolic burden imposed by cystine uptake. Finally, our proteomic and immunoblot analyses revealed reduced GPX1 abundance following SEPHS2 loss. Given the reported role of GPX1 in limiting lipid peroxidation through its interaction with OSBPL8 ^29^, its reduced abundance may further compromise antioxidant defense, although we do not have evidence of its contribution to the phenotype observed here.

In summary, our findings link impaired selenoprotein biosynthesis to ferroptotic sensitivity and cystine-associated disulfide stress. By connecting selenium utilization to cystine handling, mitochondrial function, and metabolic adaptation, these findings provide a broader framework for understanding how tumor cells maintain redox homeostasis and may reveal exploitable vulnerabilities arising from impaired redox resilience.

## Acknowledgments

J.P.F.A. acknowledges the support of from Rudolf Virchow Center, University of Würzburg and additional grants from the Deutsche Forschungsgemeinschaft (DFG), FR 3746/5-1, FR 3746/6-1, CRC/TRR205, INST269/886-1, and TRR 387/1 (514894665); the EU-H2020 (ERC-Consolidator, DeciFERR); and the Deutsche Jose Carreras Leukämie Stiftung (DJCLS 01 R/2022). H.M. is supported by a fellowship from Alexander von Homboldt Foundation. A.S and F.C.E.V. acknowledges the support from the Deutsche Forschungsgemeinschaft (DFG) grant SPP-2306 and SCHU2670/2-1/2. F.C.E.V. is supported by Excellence through Focus’ funded by zukunft.niedersachsen, the joint science funding program of the Lower Saxony Ministry of Science and Culture and the Volkswagen Foundation, UMG2032. H.A acknowledges DFG Priority Program SPP 2306 (AL 1533/5-1) and the Deutsche Jose Carreras Leukämie Stiftung (DJCLS 04 R/2024). H.G.A acknowledges DFG grant (project C05 within the CRC1366 “Vascular control of organ function” (394046768)) and ERC AdG “Angiomature” (787181). A.K.J acknowledges the support of Emmy Noether Grant from DFG, JA3274-1/1. H.A and M.E also acknowledge the German Cancer Research Center (DKFZ). H.M. and M.E. received funding from the SPP 2306 start-up funding scheme (461507177). Generative artificial intelligence was used to assist with language editing. The authors are responsible for the accuracy and integrity of the final manuscript.

## Author Contributions

H.M. performed most of the experiments and analyzed data. M.E. performed the in vivo experiments and analyzed experimental and bioinformatics data. F.P.F, H.A., M.S., Z.C., S.S., J.M., G.F. and R.S. contributed to the development of the methodology and performed experiments. M.F. and U.S. performed the radiolabeling experiments. S.O., U.B. and T.A. contributed to targeted sulfur metabolome analysis. F.C.E.V., L.S. and A.S. performed and analyzed the tracing and metabolomics data. B.G. performed MS-based proteomics experiments. H.G.A, and A.K.J supported with reagents, and infrastructure for proteomics analysis. G.F., H.M., and J.P.F.A. wrote the manuscript. The final version received input from all authors. All authors contributed to data discussion, interpretation, revision and approved the final version of the manuscript.

## Methods

### Chemicals and reagents

Liproxstatin-1 (Lip-1; SML1414), ML210 (SML0521), RSL3 (SML2234), erastin (E7781), L-cystine (C8755), βMe (21985023), 5,5′-dithiobis(2-nitrobenzoic acid) (DTNB; D8130), tris(2-carboxyethyl)phosphine hydrochloride (TCEP; C4706), sodium selenite (S5261), selenocystine (545996) and fluorescein O,O′-diacrylate (FOdA; 98%, 570257) were obtained from Sigma-Aldrich (Merck KGaA). BODIPY FL L-cystine (HY-D1605; CAS 1180490-57-6) was purchased from MedChemExpress. C11-BODIPY 581/591 (D3861) and buthionine sulfoximine (BSO; 235520010) were obtained from Thermo Fisher Scientific. 2-(N-morpholino)ethanesulfonic acid (MES; 4256.3) was obtained from Carl Roth.

Where appropriate, stock solutions of compounds used for cellular treatments were prepared in DMSO under sterile conditions and stored at −20 °C. Other reagents and fluorescent probes were prepared and stored according to the manufacturers’ instructions.

### Cell culture

The human fibrosarcoma cell line HT1080 and the human neuroblastoma cell line SK-N-DZ were obtained from the American Type Culture Collection (ATCC; CCL-121 and CRL-2149, respectively). Cells were cultured in Dulbecco’s modified Eagle’s medium (DMEM; Gibco) supplemented with 10% fetal bovine serum (FBS; Gibco) and penicillin–streptomycin (100 U ml⁻¹; Gibco) at 37 °C in a humidified atmosphere containing 5% CO_2_. Cell lines were routinely tested for mycoplasma contamination. SEPHS2-KO cells were maintained in complete culture medium supplemented with Lip-1 (500 nM) and erastin (250 nM).

### Cell viability and clonogenic growth assays

Cell viability was assessed using the Alamar Blue assay. For experiments involving SEPHS2-KO cells, 2,000 cells were seeded per well in 96-well plates and pretreated with Lip-1 (500 nM) for 6 h, unless otherwise stated. The indicated concentrations of BSO, βMe, cystine, RSL3 or erastin were then added, and cells were incubated for 48 h.

Resazurin sodium salt (1 g) was dissolved in 200 ml sterile PBS, and the solution was filtered through a 0.22-µm filter and stored at 4 °C. Immediately before use, the stock solution was diluted by adding 100 µl to 10 ml growth medium. The working solution was added to the cells, followed by incubation for 2 h. Fluorescence was measured using a Spark microplate reader (Tecan, Zürich, Switzerland) with excitation and emission filters of 540/35 nm and 590/20 nm, respectively. For the clonogenic assay, 15,000 cells were seeded per well in 12-well plates. The indicated treatments were initiated 6 h after seeding, and cells were maintained under the respective experimental conditions for 6 days. Colonies were subsequently stained with 0.01% (w/v) crystal violet in PBS.

### Generation of knockout cell lines

Single-guide RNAs (sgRNAs) targeting the genes of interest were selected using the Vienna Bioactivity CRISPR (VBC) score (https://vbc-score.org/). Corresponding oligonucleotides were designed with overhangs compatible with BsmBI-digested sites (Microsynth) and cloned into the pLentiCRISPR-v2 vector. Constructs were verified by Sanger sequencing.

Lentiviral particles were produced in HEK293T cells using a third-generation packaging system. Briefly, HEK293T cells were transfected with the lentiviral transfer plasmid, the packaging plasmids pMDLg/pRRE (Addgene, #12251) and pRSV-Rev (Addgene, #12253), and the envelope plasmid pCMV-VSV-G (Addgene, #8454), using X-tremeGENE HP DNA Transfection Reagent (Sigma-Aldrich, 6366546001). Viral supernatants were collected 48 h after transfection and filtered through a 0.45-µm membrane. Target cells were transduced with filtered viral supernatants with protamine sulphate solution (final concentration 8µg/ml) and subsequently selected for 10 days. Selected cells were used as pooled populations or expanded as single-cell clones, as indicated. Knockout was validated by assessing loss of the corresponding protein by Western blotting.

The following sgRNAs were used in this study: SEPHS2_sgRNA1, CACCGTAGAGGTCACTCAGCACGT; Non-target_SgRNA, ATCTAGTCCTCTAATCGAT; TXNRD1_sgRNA1, GAAGCTGTACAGAATCACAT; TXNRD1_sgRNA2, GATAGGATGCTCCAACAACCA; TXNRD1_sgRNA3, GTTACTGCCCGGGTAAGACCC; TXNRD1_sgRNA3, GTTACTGCCCGGGTAAGACCC; SECISBP2_sgRNA1, GCGGTTGAGAGCAAACACAAAGG; EEFSEC_sgRNA1 GATGATGGTCCGGATGAGGGAGG; EEFSEC_sgRNA3 GCCATGCAAGGAGACCGGCTGGG; SLC7A11_sgRNA1, CACCGA ACA AAG CTGGGATGA ACA G and NFE2L2_sgRNA, GGATCTGCCAACTACTCCC. Lentiviral production and transduction were performed in a biosafety level 2 laboratory.

### Generation of gene-overexpression constructs and stable cell lines

Codon-optimized WT SEPHS2 or TXNDC17 coding sequence was ordered as gBlock from IDT and amplified by PCR for cloning. SEPHS2 variants carrying the U60A and U60C substitutions were generated in parallel. All SEPHS2 constructs, including the wild-type and mutant variants, carried an AviTag; the TXNDC17 construct carried an ALFA tag.

These constructs were cloned into the p442-IRES backbone to yield p442-IRES-Mock (empty vector), p442-IRES-SEPHS2-AviTag, p442-IRES-SEPHS2-U60A-AviTag, p442-IRES-SEPHS2-U60C-AviTag, p442-IRES-NFE2L2, p442-IRES-SLC7A11 and p442-IRES-TXNDC17-ALFA-tag. SEPHS2 and SLC7A11 overexpression constructs were generated using both the p442-IRES lentiviral expression system and the pSB-ET-iE Sleeping Beauty transposon system, as described previously ^30^. For transposon-based expression, pSB expression vectors were co-transfected with the Sleeping Beauty transposase vector pCMV-(CAT)T7-SB100 to facilitate genomic integration and establish stable expression cell lines. All plasmids were verified by Sanger sequencing before use.

### Immunoblotting

Cell lysates were prepared in radioimmunoprecipitation assay (RIPA) buffer supplemented with a protease inhibitor cocktail (Roche). For non-reducing and corresponding reducing immunoblot analysis, cells or excised tumor tissues were lysed and homogenized in RIPA buffer supplemented with 10 mM N-ethylmaleimide to preserve protein thiol residues during sample preparation. Samples were prepared using SDS loading buffer without βME for non-reducing immunoblotting or with βME for reducing immunoblotting. Proteins were separated by SDS–polyacrylamide gel electrophoresis (SDS–PAGE) and electrotransferred onto polyvinylidene difluoride (PVDF) membranes. Membranes were blocked for 1 h at room temperature in 5% non-fat milk prepared in Tris-buffered saline containing 0.1% Tween 20 (TBS-T). Primary antibodies diluted in 5% bovine serum albumin (BSA) in TBS-T were incubated with the membranes overnight at 4 °C. After washing with TBS-T, membranes were incubated for 2 h at room temperature with horseradish peroxidase (HRP)-conjugated secondary antibodies (1:3,000; anti-rabbit, #7074; anti-mouse, #7076; or anti-rat, #7077; Cell Signaling Technology) diluted in 5% non-fat milk in TBS-T. Protein bands were visualized using Immobilon Western Chemiluminescent HRP Substrate (Sigma-Aldrich), and chemiluminescence was detected using an ImageQuant 800 system (Cytiva).

The primary antibodies used were anti-VINCULIN (1:5,000, sc-73614, Santa Cruz Biotechnology), anti-SEPHS2 (1:1,000, 14109-1-AP, Proteintech), anti-GPX4 (1:1,000, ab125066, Abcam), anti-xCT/SLC7A11 (1:10; raised against recombinant human xCT by the group of M. Conrad, Munich, Germany), anti-SELENON (1:1,000, sc-365824, Santa Cruz Biotechnology), anti-SELENOS (1:1,000, HPA010025, Sigma-Aldrich), anti-TXNDC17/TRP14 (1:1,000, 20811-1-AP, Proteintech), anti-NRF2 (1:1,000, ab62352, Abcam), anti-TXNRD1 (1:1,000, D1T3D, Cell Signaling Technology), anti-β-ACTIN (1:5,000, A5441, Sigma-Aldrich), anti-PRDX1 (1:1,000, 8499, Cell Signaling Technology), EEFSEC (1:1,000, 10628-1-AP, Proteintech) SECISBP2 (1:1,000, 12798-1-AP, Proteintech) HO-1 (1:1,000, 70081, Cell Signaling Technology) and anti-Drebrin (1:1,000, 10260-1-AP, Proteintech).

### Assessment of lipid peroxidation

SEPHS2-KO HT1080 cells were seeded at 200,000 cells per well in six-well plates one day before analysis. On the following day, culture medium containing Lip-1 and erastin was replaced with fresh medium lacking both supplements, and cells were incubated for 14 h. Lipid peroxidation was assessed using C11-BODIPY 581/591. Separate samples were incubated with C11-BODIPY 581/591 (1 µM) for 30 min at 37 °C. SEPHS2-KO cells reconstituted with SEPHS2 WT were used as the reference control. Cells were then trypsinized, resuspended in 500 µl fresh PBS and analyzed by flow cytometry using a BD FACSCanto II flow cytometer (BD Biosciences) with 488-nm excitation. Fluorescence was recorded in the FITC and PE channels, and data were analyzed using FlowJo software (BD Biosciences).

Cell populations were gated using forward-scatter area (FSC-A) and side-scatter area (SSC-A) to exclude debris and select the cell population of interest. Unstained cells were used to determine background fluorescence. Lipid peroxidation was quantified by measuring the fluorescence intensity of oxidized C11-BODIPY in the FITC channel, with increased fluorescence indicating elevated lipid peroxidation.

### Human cancer cell line datasets

Human cancer cell line datasets were obtained from the DepMap portal (https://depmap.org/portal).

### Direct labelling with radioactive selenium

For direct labelling with radioactive sodium selenite (^75^Se-labelled Na_2_SeO_3_; Polatom), SEPHS2-KO HT1080 cells were seeded in 6-cm dishes in serum-free DMEM supplemented with Lip-1 (500 nM), erastin (250 nM) and 6 mCi ^75^Se-labelled sodium selenite. After 48 h of incubation, cells were harvested and lysed in RIPA buffer. Proteins were separated by 12% SDS–PAGE, and gels were exposed to phosphor screens for one week to detect radiolabeled proteins.

### Proteomic analysis

#### Proteomic sample preparation

Cell pellets were lysed in 100 µL of 1% sodium deoxycholate (SDC) buffer, followed by heat denaturation at 95°C for 15 minutes and sonication using a Bioruptor (20 cycles of 30 s on/off) as described previously ^31^. Proteins were digested overnight at 37°C and 1,400 rpm with a Trypsin/LysC mixture (Sigma-Aldrich T6567-1MG; Fujifilm Wako 121-05063/125-05061) at a 1:100 enzyme-to-protein ratio. Digestion was quenched by addition of five volumes of isopropanol containing 1% trifluoroacetic acid (TFA). Peptides were desalted using StageTips packed with three layers of Styroldivinylbenzol-Reversed Phase Sulfonate (SDB-RPS) Empore™ SPE disks (66886-U), pre-equilibrated with 30% methanol/1% TFA and 0.2% TFA. Following sample loading (30 µg), StageTips were washed sequentially with isopropanol/1% TFA and 0.2% TFA, and peptides were eluted with 60 µL of 80% acetonitrile (ACN)/1.25% ammonium hydroxide. Eluates were dried at 45°C for 90 minutes in a vacuum concentrator and resuspended in buffer A* (2% ACN, 0.1% TFA). Peptide concentrations were determined by NanoDrop and samples were stored at −20°C until analysis.

#### Data-independent acquisition (DIA) mass spectrometry

Peptides (400ng per injection) were separated on a proteoElute liquid chromatography system (Bruker Daltonics) coupled to a timsTOF HT mass spectrometer via a CaptiveSpray nano-electrospray ion source. Chromatographic separation was performed on an IonOpticks Aurora Ultimate 25 cm × 75 µm C18 UHPLC column using a binary gradient of buffer A (0.1% formic acid, 2% ACN) and buffer B (0.1% formic acid, 90% ACN) at a flow rate of 0.3 µL/min. The gradient was as follows: 2% to 12% B over 60 minutes, 12% to 20% B over 30 minutes, 20% to 30% B over 10 minutes, 30% to 85% B over 10 minutes, followed by a 10-minute hold at 85% B. TIMS calibration was performed using Agilent ESI-Low Tuning Mix reference ions (m/z 622, 922, and 1,222) with known inverse reduced ion mobility coefficients (1/K0). Data were acquired in dia-PASEF mode over an m/z range of 100–1,700 and 1/K0 of 0.65–1.40. Collision energy was ramped linearly from 20 eV at 1/K0 = 0.6 Vs/cm^2^ to 80 eV at 1/K0 = 1.60 Vs/cm^2^. TIMS acquisition parameters were: scan range 100–1,700 m/z, ramp time 100 ms, 100% duty cycle, cycle time 100 ms, and spectral acquisition rate of 9.43 Hz.

#### MS data processing

Raw data were processed using DIA-NN (v1.8.2) ^32^ with default parameters and searched against the Homo sapiens UniProt reference proteome (accessed 7 March 2021). Trypsin/P was specified as the protease, allowing up to one missed cleavage. N-terminal methionine excision and cysteine carbamidomethylation were set as fixed modifications. A theoretical spectral library was constructed using deep learning-based prediction, and a decoy database was used to estimate the false discovery rate (FDR). FDR thresholds were set at 1% for both precursor and protein levels, with a minimum of one peptide required per protein identification.

The mass spectrometry proteomics data have been deposited to the ProteomeXchange Consortium via the PRIDE ^33^ partner repository with the dataset identifier PXD087432.

#### Proteomics of tumor tissue form in-vivo orthotopic experiment

Tumor tissue samples were collected, immediately snap-frozen in liquid nitrogen, and stored at −80°C until processing. For protein extraction, a small piece of tumor tissue was homogenized in ice-cold lysis buffer containing 8 M urea and 50 mM Tris-HCl (pH 8.0), supplemented with a protease inhibitor cocktail. Homogenization was performed under cold conditions to minimize protein degradation. Homogenates were incubated on ice for 20–30 minutes with occasional mixing and subsequently centrifuged at 16,000 × g for 15–20 minutes at 4°C to remove insoluble debris. Supernatants were transferred to fresh tubes, and total protein concentration was determined by Bradford assay. Protein extracts were stored at −80°C until further processing. LC-MS/MS acquisition and raw data processing were performed at the Proteomics Core Facility, German Cancer Research Center (DKFZ), Heidelberg, Germany.

### Bioinformatic analysis

#### Proteomic data analysis

Raw protein intensity data were processed and analyzed in R using the limma package. Proteins with zero intensity across all samples were excluded. Remaining values were log2-transformed after replacing zeros with missing values (NA). Samples were normalized by median centering, shifting each sample median to the global median. Missing values were imputed using a Perseus-style down-shift strategy, replacing missing entries with random values drawn from a normal distribution shifted 1.8 standard deviations below the sample mean with a width of 0.3 standard deviations, to model the distribution of low-abundance proteins near the detection limit (seed = 42). Differential protein expression was assessed using a linear model framework in limma. A design matrix was constructed without an intercept term and pairwise contrasts were defined between experimental and control groups. Model fitting was performed using lmFit, and empirical Bayes moderation was applied via eBayes with robust and trend corrections. Differential expression statistics were extracted using topTable without multiple testing correction. Proteins with log2 fold change > 0.5 or < −0.5 and p < 0.05 were classified as significantly up-or down-regulated, respectively. Results were visualized as volcano plots using ggplot2, with the top 10 most up-and down-regulated proteins labelled using ggrepel.

#### Functional enrichment analysis

Gene symbols were converted to Entrez IDs using clusterProfiler package (v4.18.4), with annotation provided by the org.Hs.eg.db database. Pathway enrichment analysis was performed separately for up-and down-regulated protein sets using the enricher function in clusterProfiler, with gene sets retrieved from the WikiPathways database (Homo sapiens, October 2026 release, 991 pathways). Over-representation analysis was based on a hypergeometric test, with pathway size restricted to 5–500 genes, and results were filtered at an FDR threshold of less than 0.05. The five most significantly enriched pathways from each direction were selected for visualization. For each enriched pathway, the mean log2 fold change of member proteins was calculated to provide directional context. Results were visualized as dot plots in which dot size reflects −log10(FDR) and dot color reflects mean log2 fold change, ranging from dark blue (down-regulated) to dark red (up-regulated).

### Statistical analysis of proteomics data

Statistical analysis was performed using MetaboAnalyst ^34^. Spectronaut output files containing protein identifications and label-free quantification (LFQ) intensities were used as input. Features identified as contaminants or reverse sequences were removed before further analysis. Missing values were imputed using the limit of detection, defined as one-fifth of the minimum positive value for the corresponding variable. The data were then log_10_-transformed for statistical analysis. Differentially expressed proteins (DEPs) were defined using a fold-change threshold of 1.5 and a raw P value cutoff of 0.05. Volcano plots and heatmaps displaying the top DEPs were generated in MetaboAnalyst. Metascape was used to identify statistically enriched terms and pathways among the DEPs, including Gene Ontology (GO) terms, Kyoto Encyclopedia of Genes and Genomes (KEGG) pathways and canonical pathways. Analyses were performed using five replicates.

### Measurement of SLC7A11 transport activity

SLC7A11 transport activity was assessed using seleno-L-cystine as an L-cystine analogue and fluorescein O,O′-diacrylate (FOdA) as a fluorophore that reacts with L-selenocysteine, as described by ^35^. Cells were seeded in six-well plates 24 h before the experiment and maintained under standard culture conditions. On the day of the experiment, cells were incubated in cystine-free medium (Gibco, cat. no. 21013024) supplemented with 200 µM seleno-L-cystine, dissolved in 1 M HCl, for 30 min at 37 °C in a humidified incubator containing 5% CO_2_. Substrate uptake was stopped by washing the cells with ice-cold PBS, followed by extraction of soluble metabolites with ice-cold methanol.

Cell extracts were subsequently incubated in 100 mM MES buffer (pH 6.0) containing 10 µM FOdA and 200 µM TCEP for 30 min at 37 °C to reduce seleno-L-cystine to L-selenocysteine. The supernatant was transferred to a black 96-well plate, and fluorescence was measured at excitation and emission wavelengths of 485 and 535 nm, respectively, using a Spark microplate reader (Tecan). For normalization, the cells remaining in the six-well plates were washed with PBS and lysed in 0.5 M NaOH overnight. Protein concentrations were determined using the Pierce 660 nm Protein Assay Kit (Gibco, cat. no. 22662). Fluorescence intensities were normalized to total protein content and expressed as relative SLC7A11 transport activity.

### TXNRD1 activity assay

TXNRD1 activity was assessed using a 5,5′-dithiobis(2-nitrobenzoic acid) (DTNB) reduction assay. Cell lysates (2.5 mg/mL) were preincubated with DMSO or the TXNRD1 inhibitor TRi-1 (1 µM) for 20 min in the presence of 250 µM NADPH. For the assay, 20 µl of each lysate was added to a reaction mixture containing 2.5 mM DTNB and 250 µM NADPH in 0.2 M phosphate-buffered saline (PBS) supplemented with 10 mM EDTA. Absorbance at 412 nm was monitored for 30 min. TXNRD1 activity was estimated from the difference in DTNB reduction between DMSO-treated and TRi-1-treated samples.

### Confocal microscopy of actin filaments

Cells were seeded in imaging chambers (Cellvis) and cultured for 24 h before treatment with 200 µM BSO for 6 h. After treatment, cells were washed three times with PBS and fixed with 4% formaldehyde in PBS for 15 min at room temperature. Residual aldehydes were quenched with 0.1 M glycine in PBS for 10 min, followed by three washes with PBS. Cells were permeabilized with 0.5% Triton X-100 in PBS for 10 min, washed three times with PBS and blocked with 3% donkey serum in PBS for 10 min at room temperature. F-actin was stained with Alexa Fluor 647-conjugated phalloidin (40 nM) for 30 min at room temperature in the dark. Cells were subsequently washed three times with PBS, counterstained with DAPI (1 µg/ml) for 5 min and washed again three times with PBS. Confocal images were acquired using a Leica SP8 confocal microscope equipped with a HCX PL APO CS 63×/1.4 NA oil-immersion objective.

### Analysis of oxygen consumption rate

Oxygen consumption rate (OCR) and extracellular acidification rate (ECAR) were measured in SK-N-DZ SEPHS2-KO cells transduced with the mock or DOX-inducible SEPHS2 WT construct under the following conditions: in the presence of liproxstatin-1, with or without erastin, and with or without DOX. Measurements were performed using the Seahorse XF96 Extracellular Flux Analyzer (Agilent Technologies) in conjunction with the Seahorse XF Cell Mito Stress Test Kit.

Sequential injections of oligomycin (2.5 µM), carbonyl cyanide-4-(trifluoromethoxy) phenylhydrazone (FCCP; 2 µM), and rotenone (0.5 µM) were administered during the assay according to the manufacturer’s protocol.

For assay preparation, cells were expanded in T25 flasks under standard culture conditions. Prior to the experiment, cells were detached using Accutase, washed twice with Seahorse XF RPMI assay medium (pH 7.4, supplemented with 10 mM glucose, 1 mM pyruvate, and 2 mM glutamine), and seeded at a density of 2 × 10^5^ cells per well. Cells were allowed to equilibrate for one hour at 37°C in a non-CO_2_ incubator prior to measurement.

### LC-MS Metabolomics

For intracellular metabolites, cells were washed with cold saline solution (B.Braun # 3570130), snap frozen in liquid nitrogen and scraped off in 500µl ice-cold MeOH/H2O/Acetonitrile (50/20/30 v/v) containing internal standards (DLM-372-PK and DLM-3106-5, Cambridge Isotope laboratories). For stable isotope tracing experiments, labelled metabolites for Glucose (^13^C_6_; #CLM-1396-1) or L-Glutamine (^13^C_5_; #CNLM-1822-H) were purchased from Cambridge Isotope Laboratories. Tomur material from SLC7A11-overexpressing SK-N-DZ cells were extracted using Blight & Dyer protocol ^36^. Frozen tumors samples were homogenized using the retsch-mill in ice-cold extraction buffer to quench metabolism. Around 40mg of tumor material was extracted and volumes of buffers were adjusted accordingly to normalize differences in wight. Upper, polar phase from Bligh & Dyer extraction were transferred into new tubes and dried in a SpeedVac before further reconstitution for LCMS analysis. For stable isotope tracing experiments on SK-N-DZ cells overexpressing SLC7A11, labelling time was performed for 24h using standard medium conditions with dialyzed serum. Metabolite extraction of tumor cells after stable isotope tracing was performed by solid phase extraction (SPE) following standard protocols.

Dried metabolite extracts were dissolved in 100 μl 5 mM NH4OAc in CH3CN/H2O (75:25, v/v), and 3 µl of each sample was applied to an amide-HILIC column (2.6 μm, 2.1×100 mm, Thermo Fisher, 16726-012105). Metabolites were separated at 30 °C by LC using a DIONEX Ultimate 3000 UPLC system and the following solvents: solvent A consisting of 5 mM NH4OAc in CH3CN/H2O (5:95, v/v) and solvent B consisting of 5 mM NH4OAc in CH3CN/H2O (95:5, v/v). The LC gradient program was: 98% solvent B for 2 min, followed by a linear decrease to 40% solvent B within 5 min, then maintaining 40% solvent B for 13 min, then returning to 98% solvent B in 1 min and then maintaining 98% solvent B for 5 min for column equilibration before each injection. The flow rate was maintained at 350 μl/min. The eluent was directed to the hESI source of the Q Exactive mass spectrometer (QE-MS; Thermo Fisher Scientific) from 1.85 min to 18.0 min after sample injection. The scan range was set to 69.0–1000 m/z with a resolution of 70,000 and polarity switching (negative and positive ionization). Peaks corresponding to the calculated metabolites masses taken from an in-house metabolite library (MIM +/− H+ ± 2 mmU) were integrated using the El-MAVEN software v.12.1-beta ^37^. Data from stable isotope tracing were normalized towards internal standard and cell count; data from tumor metabolomics were normalized towards internal standard and summed metabolites signal intensity. The R package IsoCorrectoR ^38^ was used to correct for natural isotope abundance.

### Targeted sulfur metabolomics analysis by LC-MS/MS

HT1080 SEPHS2-KO cells reconstituted with SEPHS2 WT or the mock control were seeded at a density of 0.25 × 10^6^ cells per well in 6-well plates and treated with DMSO (vehicle) or 200 µM BSO for 6 h. Following treatment, cells were washed twice with ice-cold PBS, and the plates were snap-frozen by placing their bottoms in liquid nitrogen and stored at −80 °C until further processing.

For metabolite extraction, plates were thawed, and 500 µL of extraction buffer containing 5 mM HPE-IAM, 70% methanol and 50 mM sodium acetate (pH 6.5) were added to each well. Cells were scraped, collected into tubes and disrupted by probe sonication at 30% intensity for 10 s, repeated twice. The lysates were incubated at 37 °C for 20 min, followed by centrifugation at 10,000 × g for 10 min at 4 °C. Subsequently, 50 µL of the supernatant was collected and mixed with 100 µL of 0.1% formic acid. The remaining protein pellets were air-dried, dissolved in 300 µL of 0.2 M NaOH containing 0.1% SDS and incubated at 60 °C for 30 min. Protein concentrations were determined using a BCA protein assay and used for normalization of metabolite levels. For LC–MS/MS analysis, 10 µL of the diluted sample was mixed with 10 µL of 50 nM isotope-labelled internal standards, and 10 µL of the resulting mixture was injected.

Metabolites were analyzed by liquid chromatography–electrospray ionization–tandem mass spectrometry (LC-ESI-MS/MS) using an LCMS-8060RX mass spectrometer (Shimadzu) coupled to a Nexera UHPLC system (Shimadzu). Chromatographic separation was performed using a YMC-Triart C18 column (50 × 2.0 mm inner diameter) with mobile phase A consisting of 0.1% formic acid in water and mobile phase B consisting of 0.1% formic acid in methanol. A linear gradient from 3% to 90% mobile phase B was applied over 15 min at a flow rate of 0.2 mL/min, with the column temperature maintained at 40 °C. The electrospray ionization probe, desolvation line and heat block temperatures were set to 300, 250 and 400 °C, respectively. The nebulizing, heating and drying nitrogen gas flow rates were maintained at 3, 10 and 10 L/min, respectively. Metabolites were identified and quantified using multiple reaction monitoring (MRM) with isotope-labelled internal standards. The MRM parameters for individual metabolites and the synthesis of isotope-labelled internal standards were described previously ^39, 40^.

### In vivo orthotopic mouse experiments

All animal studies and experimental protocols were conducted in accordance with the guidelines of the German Cancer Research Center (DKFZ) and approved by the governmental review board of the state of Baden-Württemberg, Karlsruhe District Council, under authorization number G-25/53, in compliance with German animal welfare regulations. NOD-Prkdcscid IL2rgtm1/Bcgen (NXG) female mice aged 3–4 months (Janvier Labs) were used for all experiments. Animals were housed in individually ventilated cages under controlled temperature and humidity conditions, with enriched environments including appropriate bedding material.

To establish orthotopic mouse models of neuroblastoma, 2 × 10^5^ SK-N-DZ cells were surgically transplanted into the right adrenal gland. Prior to injection, cells were resuspended in a 1:1 (vol/vol) mixture of growth factor-reduced Matrigel (Corning) and phosphate-buffered saline (PBS). A total volume of 20 µL of the cell suspension was injected into the right adrenal gland of anesthetized mice. Tumor engraftment and growth were monitored longitudinally by bioluminescence imaging using an IVIS Spectrum Xenogen device (Caliper Life Sciences). A reproducible bioluminescent signal was detected as early as five days following transplantation, confirming successful tumor establishment.

From the day of tumor cell injection, mice received doxycycline-supplemented drinking water at a concentration of 2 mg/mL to support inducible expression of SEPHS2. On day five post-injection, doxycycline was withdrawn from the KO group to promote loss of SEPHS2. Animal health and welfare were monitored daily throughout the experiment. Mice were euthanized upon reaching humane endpoints as specified in the approved experimental protocol. Animals that required early euthanasia prior to the predefined endpoint due to excessive weight loss or other termination criteria were excluded from all downstream analyses. All animal experiments, including tumor assessment and follow-up evaluations, were performed under blinded conditions, with the exception of the treatment procedures.

Similarly, to investigate the effect of SLC7A11 overexpression on neuroblastoma growth in vivo, a separate cohort of mice was orthotopically transplanted with SLC7A11-overexpressing SK-N-DZ cells using the procedure described above. Tumor progression was monitored longitudinally by bioluminescence imaging to assess the impact of SLC7A11 overexpression on tumor growth. Upon sacrifice, tumors were carefully dissected, weighed, and measured for size. Tumor samples were snap-frozen in liquid nitrogen and subsequently processed for western blot, proteomics and metabolomics.

### Quantification and statistical analysis

Details of the statistical analyses used for individual experiments are provided in the corresponding figure legends. Unless otherwise stated, data are presented as mean ± standard deviation (SD) from at least three independent biological replicates. Statistical significance was assessed using ordinary one-way ANOVA, two-way ANOVA or by unpaired two-tailed Student’s t-test or the Mann–Whitney test, as appropriate for the respective dataset. Analyses were performed using GraphPad Prism version 11.0 (GraphPad Software). Statistical significance was defined as follows: ns, not significant; P < 0.05; P < 0.01; P < 0.001; and P < 0.0001.

Sample sizes were not predetermined using statistical methods. Experiments were not randomized, and investigators were not blinded to experimental group allocation during experimentation or outcome assessment.

## Extended Data figures

**Extended Data Fig. S1.**
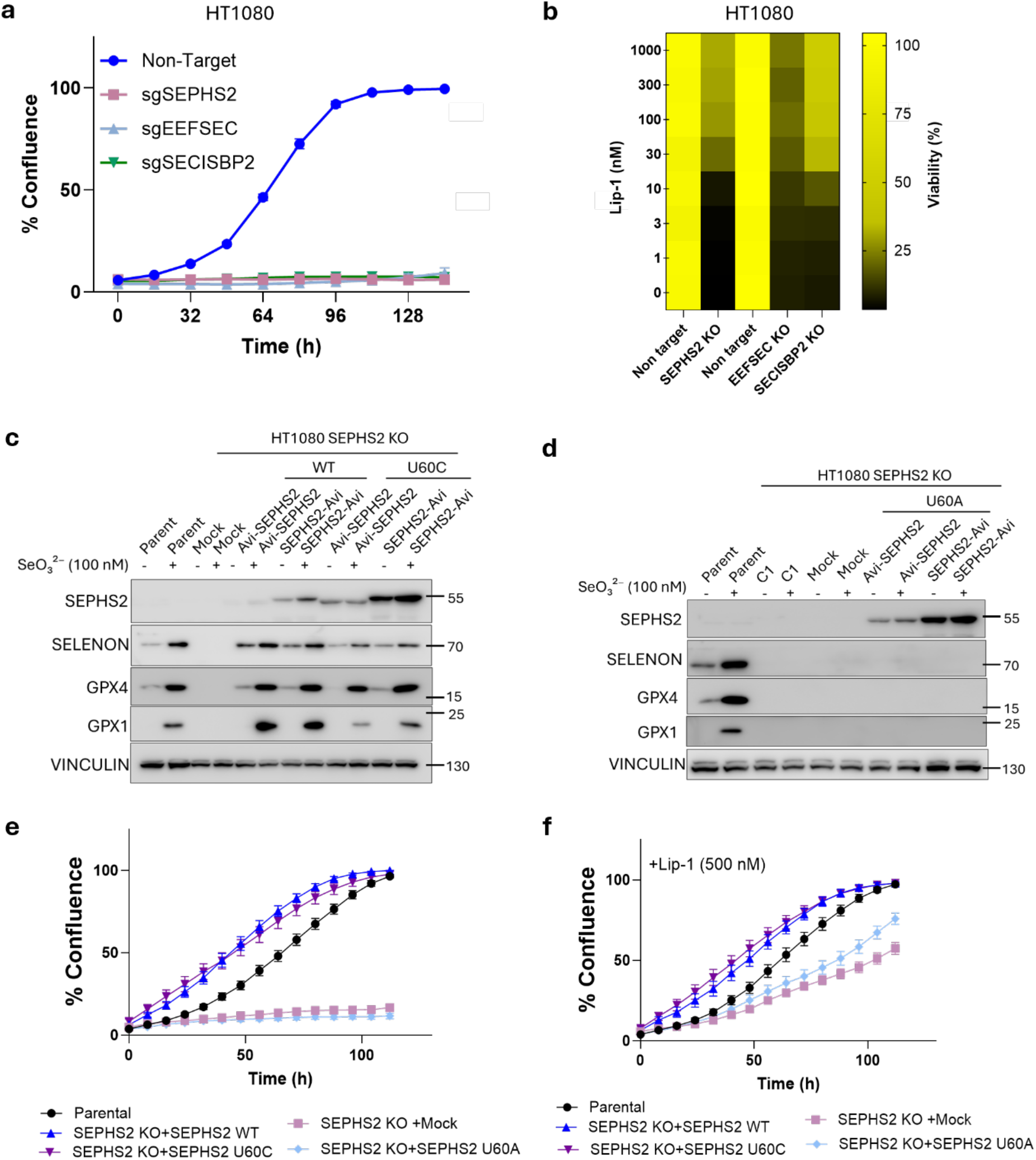
Characterization of the selenoprotein biosynthesis pathway and functional analysis of SEPHS2-deficient and sufficient cells. a, Proliferation analysis of HT1080 cells transduced with nontarget sgRNA or sgRNAs targeting SEPHS2, EEFSEC or SECISBP2. Data plotted are mean ± s.d. of triplicate wells. **b,** Viability of HT1080 cells transduced with non-target or sgRNAs targeting the indicated factors of the selenoprotein biosynthesis pathway in the presence of increasing concentrations of the ferroptosis inhibitor Lip-1. Data plotted are mean ± s.d. of triplicate wells. **c,** Immunoblot analysis of parental or SEPHS2-KO HT1080 cells transduced with the WT or U60C SEPHS2 construct, in presence or absence of 100 nM selenite. **d,** Immunoblot analysis of parental or SEPHS2-KO HT1080 cells transduced with the WT or U60A SEPHS2 construct, in presence or absence of 100 nM selenite. **e-f**, Cell proliferation analysis of SEPHS2-KO HT1080 cells reconstituted with SEPHS2 WT, U60C, or U60A mutants without (**e**) or with Lip-1 (**f**). Data plotted are mean ± s.d. of triplicate wells.

**Extended Data Fig. 2.**
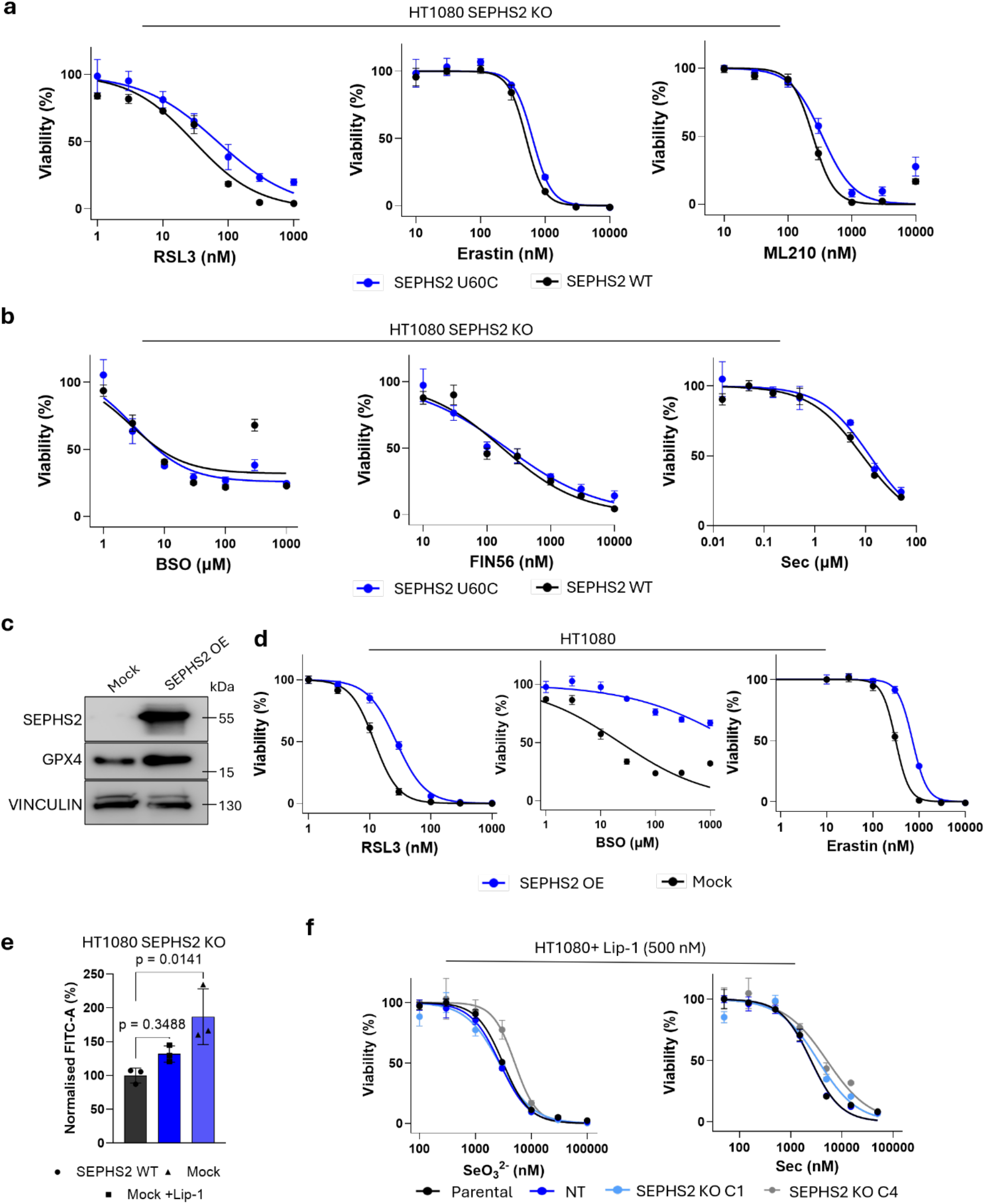
Sensitivity of SEPHS2-deficient cells reconstituted with SEPSH2 variants to ferroptosis-inducers. a, Viability of SEPHS2-KO cells reconstituted with SEPHS2 WT or SEPHS2 U60C constructs, treated with increasing concentrations of RSL3 (left) erastin (middle), and ML210 (right). Data plotted are mean ± s.d. of triplicate wells. **b,** Viability of SEPHS2-KO cells reconstituted with SEPHS2 WT or SEPHS2 U60C constructs, treated with increasing concentrations of BSO (left), FIN56 (middle) or Sec (right). Data plotted are mean ± s.d. of triplicate wells. **c,** Viability of parental HT1080 cells, transduced with non-target sgRNA or two SEPHS2 KO clones (C1 and C4), treated with increasing concentrations of selenite or Sec. Data plotted are mean ± s.d. of triplicate wells. **d,** Immunoblot analysis of SEPHS2 and GPX4 in mock and SEPHS2 overexpressing HT1080 cells. **e,** Viability of parental or SEPHS2 overexpressing cells treated with increasing concentrations of RSL3 (left), BSO (middle) or erastin (right) Data plotted are mean ± s.d. of triplicate wells. **f,** Quantification of lipid peroxidation using C11-BODIPY 581/591 staining in WT or SEPHS2-KO cells, with or without treatment with 500nM Lip-1. Data plotted are mean ± s.d. of triplicate wells.

**Extended Data Fig. 3.**
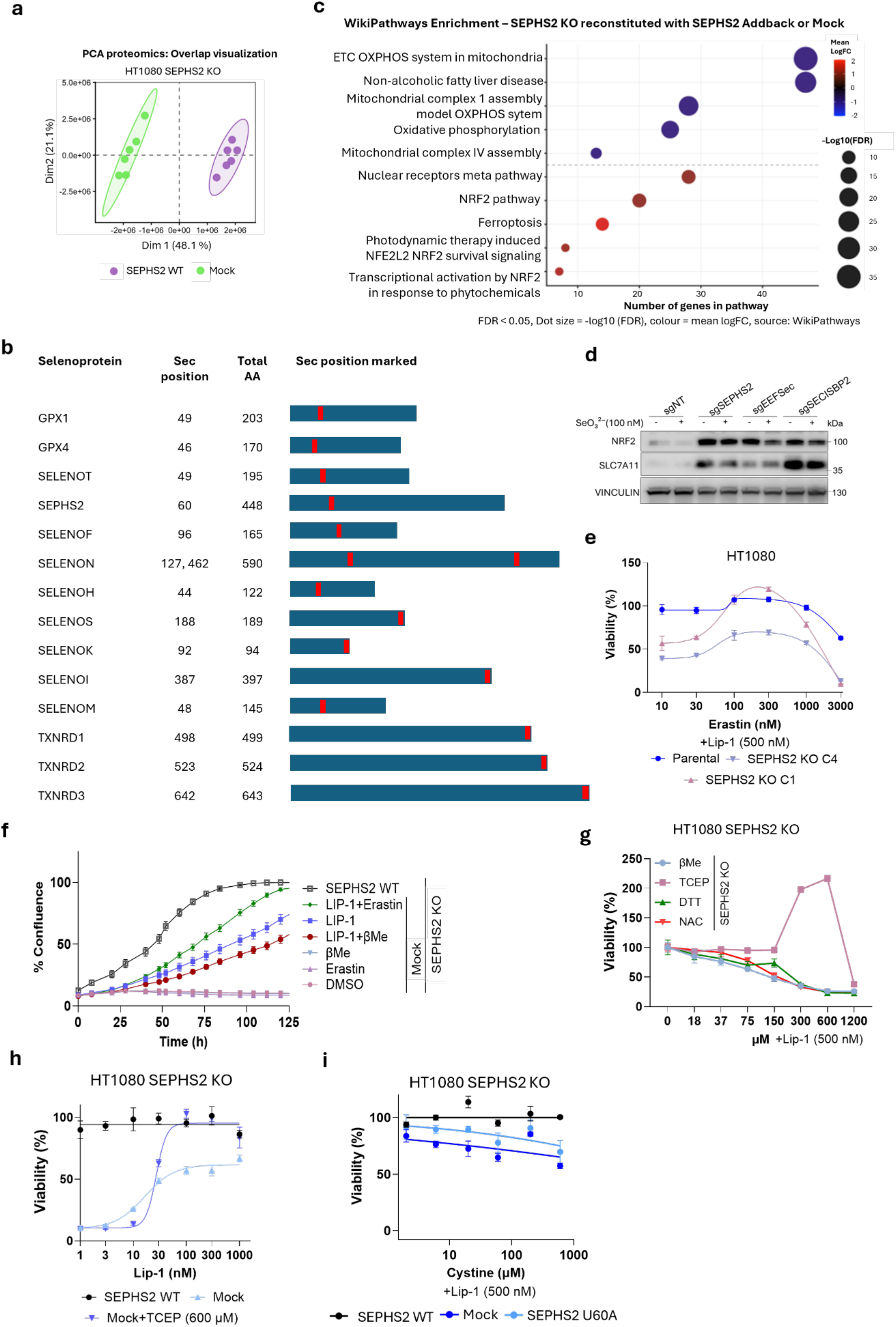
Characterization of SEPHS2-KO cells and their sensitivity to cystine-associated toxicity. a, Principal component analysis (PCA) of SEPHS2-KO HT1080 cells transduced with mock or SEPHS2 WT construct. Each dot represents one biological replicate. Dim1 and Dim2 account for 48.1% and 21.1% of the total variance, respectively. Ellipses indicate 95% confidence intervals. **b,** Schematic representation of human selenoproteins identified by proteomic analysis, showing their lengths and Sec positions (red bar). **c,** WikiPathways enrichment analysis of differentially expressed proteins between SEPHS2-KO HT1080 cells transduced with SEPHS2 WT or mock control. **d,** Immunoblot analysis of NRF2 and SLC7A11 in non-targeting control, SEPHS2-KO, EEFSEC-KO and SECISBP2-KO HT1080 cells. **e,** Viability of parental and SEPHS2 KO clones (C1 and C4), treated with increasing concentrations of erastin in the presence of Lip-1. Data plotted are mean ± s.d. of triplicate wells. **f,** Proliferation analysis of SEPHS2 KO cells treated with DMSO control, 500 nM of Lip-1, 250 nM of erastin, 50 µM of βME, or their combinations. Parental HT1080 cells are included as controls. Data plotted are mean ± s.d. of triplicate wells. **g,** Viability of SEPHS2 KO cells, treated with increasing concentrations of βME, TCEP, DTT or NAC, in the presence of 500 nM Lip-1. Data plotted are mean ± s.d. of triplicate wells. **h,** Viability of SEPHS2 KO cells transduced with mock or WT SEPHS2, treated with increasing concentrations of Lip-1, with or without 600 µM of TCEP. Data plotted are mean ± s.d. of triplicate wells. **i,** Viability of SEPHS2 KO cells transduced with mock, WT or U60A mutant constructs, treated with increasing concentrations of cystine. Data plotted are mean ± s.d. of triplicate wells.

**Extended Data Fig. 4.**
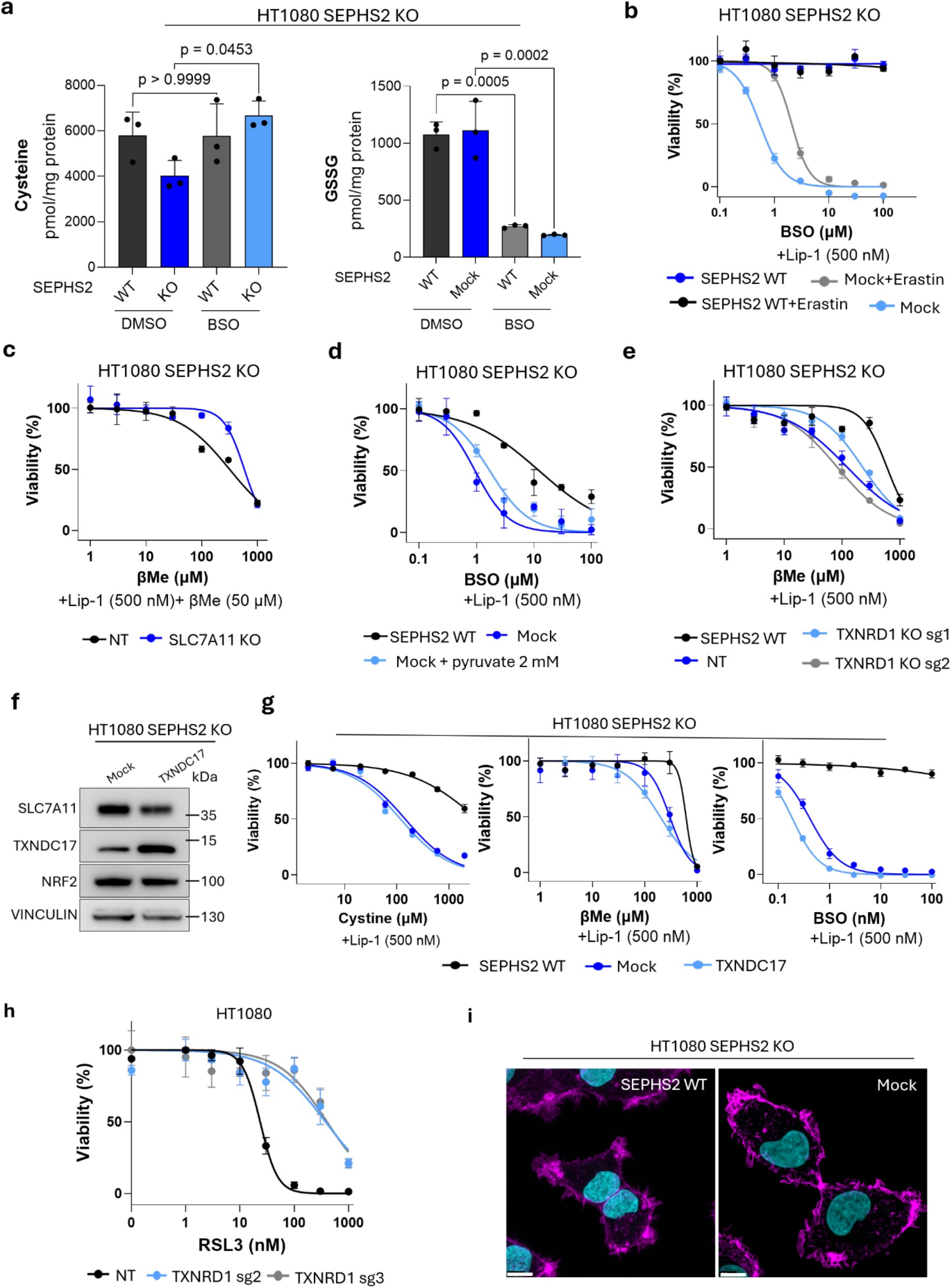
Characterization of SEPHS2-KO HT1080 cells overexpressing TXNDC17 and TXNRD1-KO HT1080 cells. a, Quantification of intracellular cysteine levels and oxidized GSH in SEPHS2-KO HT1080 cells reconstituted with SEPHS2 WT or the mock control, with or without 1μM BSO treatment, using targeted LC-MS/MS. Data plotted are mean ± s.d. of triplicate wells **b,** Viability of SEPHS2-KO HT1080 cells reconstituted with SEPHS2 WT or the mock control in response to increasing concentrations of BSO, with or without co-treatment with 250 nM erastin. Data plotted are mean ± s.d. of triplicate wells. **c,** Viability of SEPHS2 KO and SEPHS2/SLC7A11 DKO cells treated with increasing concentrations of βME. Data plotted are mean ± s.d. of triplicate wells. **d,** Viability of SEPHS2 KO cells transduced with mock or WT SEPHS2, treated with increasing concentrations of BSO in the presence or absence of pyruvate. Data plotted are mean ± s.d. of triplicate wells. **e,** Viability of SEPHS2 KO cells transduced with mock or WT SEPHS2, along with two sgRNAs targeting TXNRD1, treated with increasing concentrations of βME. Data plotted are mean ± s.d. of triplicate wells. **f,** Immunoblot analysis of SEPHS2 KO HT1080 cells, transduced with a mock or TXNDC17 overexpressing construct. **g,** Viability of cells from **f**, treated with increasing concentrations of cystine (left), βME (middle) or BSO (right). **h,** Viability of HT1080 cells transduced with non-target or two different sgRNAs targeting TXNRD1. In the presence of increasing concentrations of RSL3. Data plotted are mean ± s.d. of triplicate wells. **i,** Phalloidin staining (magenta) for F-ACTIN in SEPHS2 KO HT1080 cells transduced with a mock or SEPHS2 WT overexpressing construct. DNA is counterstained with DAPI (cyan). Scale bar is 10 μm.

**Extended Data Fig. 5.**
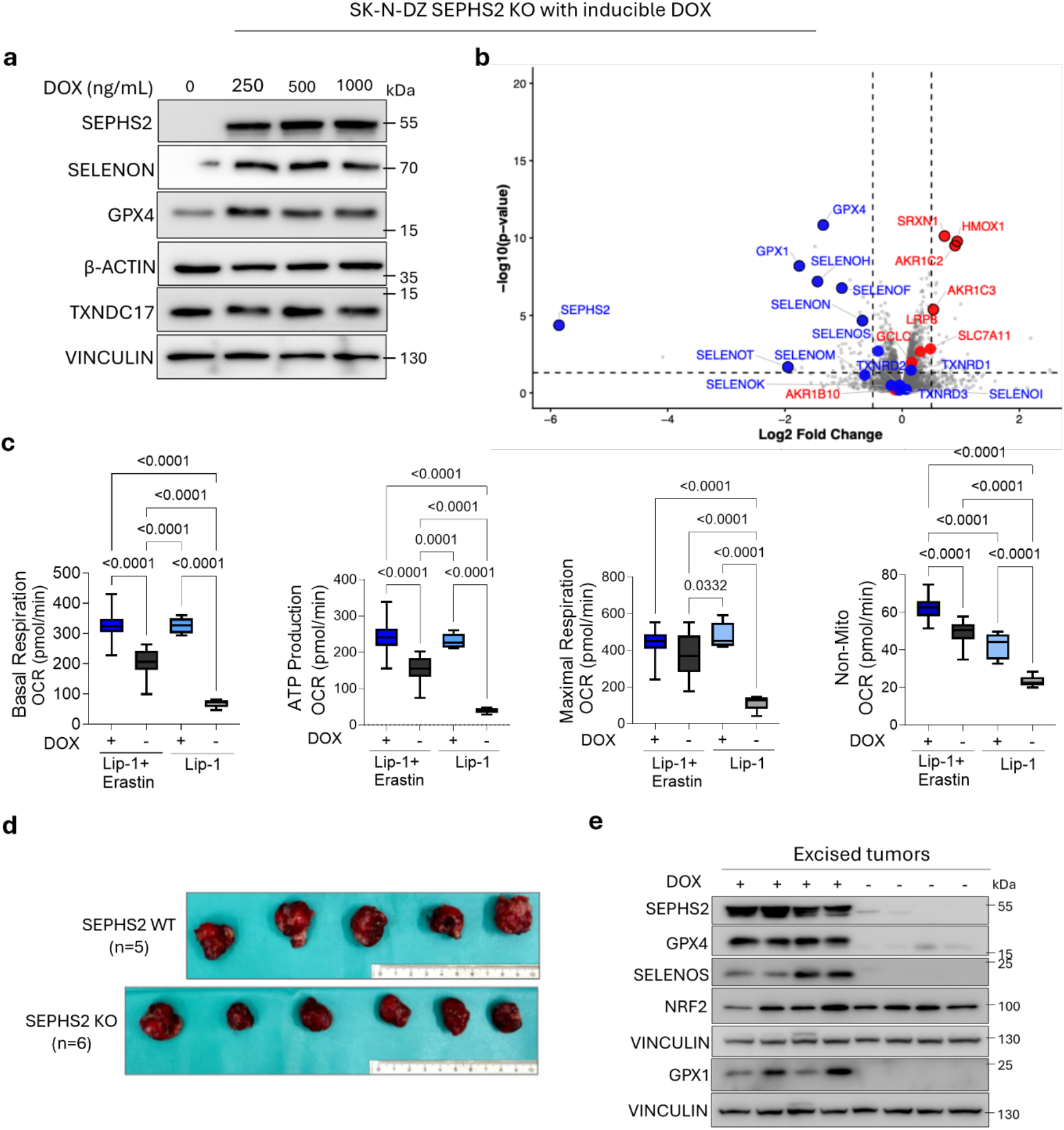
Characterization of DOX inducible SEPHS2 expression in SEPHS2-KO SK-N-DZ cells and tumor samples. a, Immunoblot analysis of SEPHS2-KO SK-N-DZ cells, reconstituted with DOX-inducible SEPHS2 WT construct, treated with increasing concentrations of DOX for 48 h. **b,** Proteomic analysis of SEPHS2-KO SK-N-DZ cells reconstituted with DOX-inducible SEPHS2 WT construct, with or without 500 ng/ml DOX treatment. Detected selenoproteins are shown in blue. NRF2 target genes are shown in red. **c,** Mitochondrial OCR analysis of SEPHS2-KO SK-N-DZ cells, reconstituted with DOX-inducible SEPHS2 WT construct, with or without DOX treatment. Data plotted are mean ± s.d. of n=5 repeats. **d,** Representative images of excised tumors on day 28 post injection with SEPHS2-KO SK-N-DZ cells reconstituted with DOX-inducible SEPHS2 WT construct, with or without 2mg/mL DOX treatment in the drinking water. **e**, Immunoblot analysis of excised tumor lysates collected from mice with or without DOX treatment (2 mg/mL) to induce SEPHS2 expression.

**Extended Data Fig. 6.**
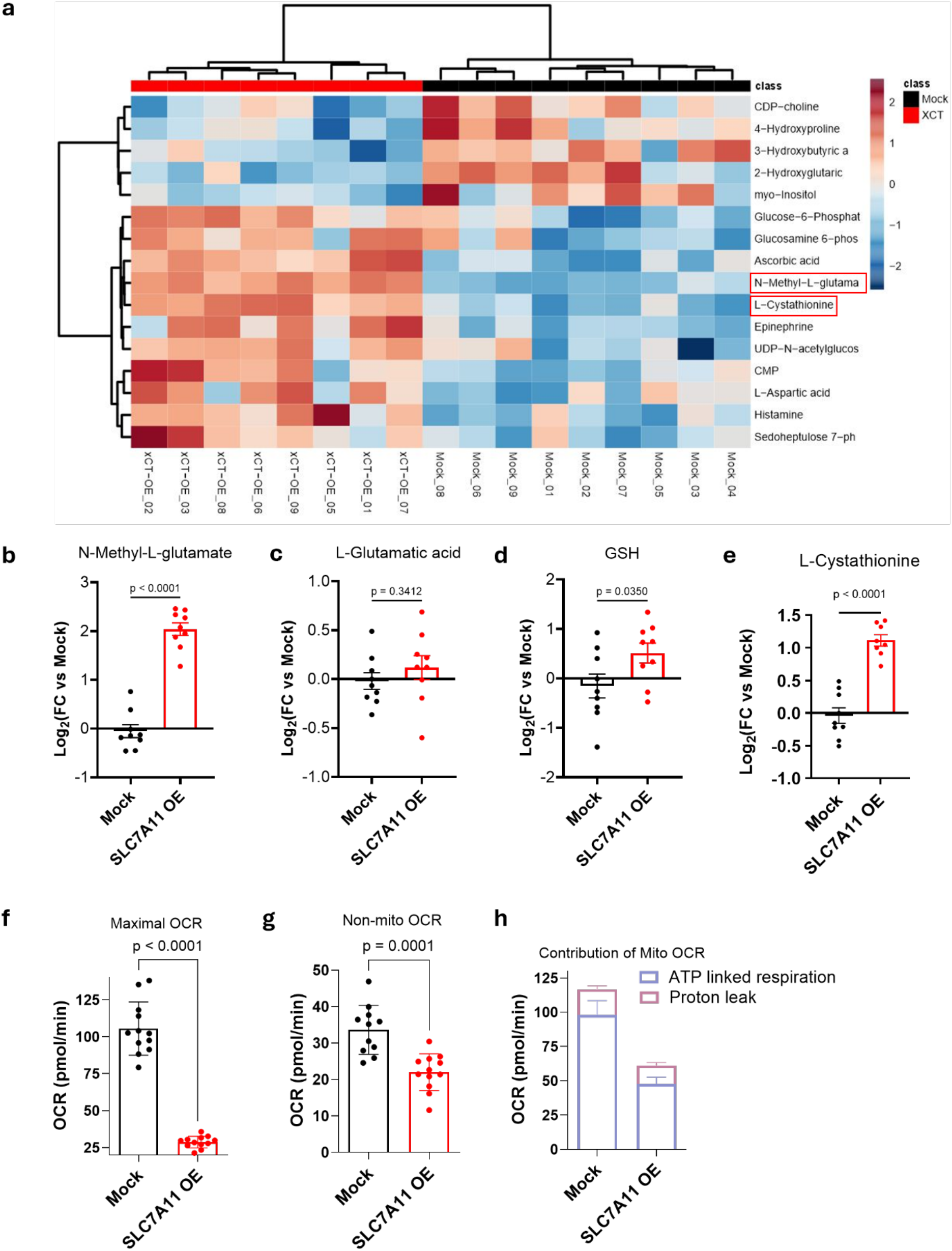
Characterization of SLC7A11 overexpressing SK-N-DZ neuroblastoma tumors. a, Heatmap showing the relative abundance of the indicated metabolites in mock and SLC7A11 overexpressing tumors. **b-e,** Relative abundance of N-methyl-L-glutamate (**b**), L-glutamic acid (**c**) GSH (**d**) or cystathione (**e**) in mock and SLC7A11 overexpressing tumors. Data plotted are mean ± s.d. of n= 8 tumors from each group. **f-h**, Parameters of oxygen consumption rate analysis of mock and SLC7A11 overexpressing cells showing maximal respiration (**f**), non-mitochondrial respiration (**g**), and ATP-linked respiration (**h**). Data plotted are mean ± s.d. of n= 10 repeats.

**Extended Data Fig. 7.**
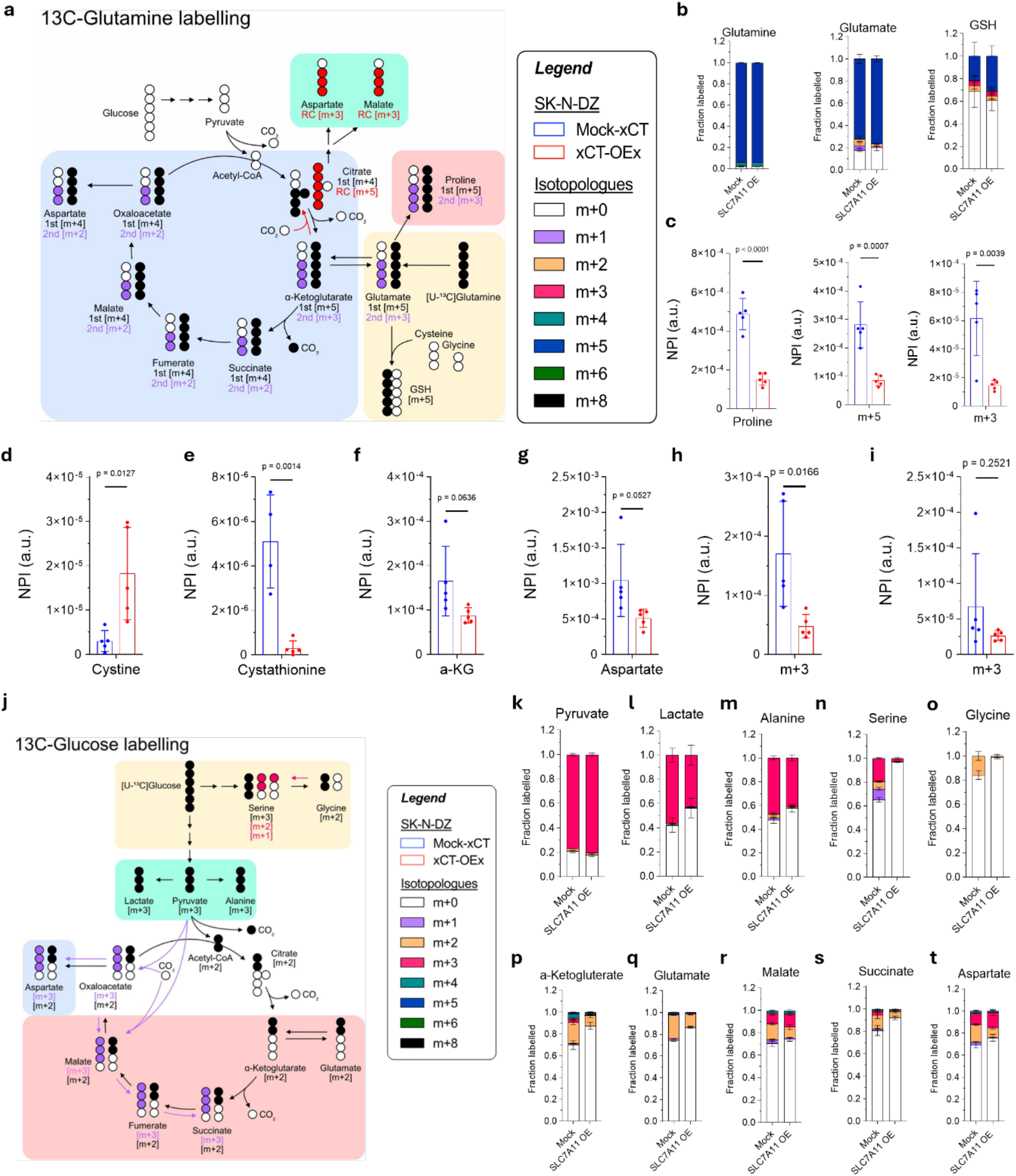
SLC7A11-overexpression in SK-N-DZ cells reduces TCA activity and changes central carbon metabolism. a, Schematic representation of ^13^C-glutamine tracing and the corresponding isotopologues in central carbon and amino acid metabolism. **b,** Fractional isotopologue distributions of glutamine, glutamate and GSH in mock and SLC7A11 overexpressing cells following 24 h of ^13^C-glutamine labelling Data plotted are mean ± s.d. of n= 5 repeats. **c,** Relative abundance of proline isotopologues (NPI: Normalized peak intensity) (total, m+5 and m+3). Data plotted are mean ± s.d. of n=5. **d-g,** Relative abundance of cystine (**d**), cystathionine (**e**), α-ketoglutarate (**f**) and aspartate (**g**). Data plotted are mean ± s.d. of n=5 repeats. **h-i,** Relative abundance of the m+3 isotopologues of aspartate (**h**) and malate (**i**). Data plotted are mean ± s.d. of n=5 wells. **j,** Schematic representation of ^13^C-glucose tracing and the corresponding isotopologues in central carbon and amino acid metabolism. **k-t**) Fractional isotopologue distributions of pyruvate (**k**), lactate (**l**), alanine (**m**), serine (**n**), glycine (**o**), α-ketoglutarate (**p**), glutamate (**q**), malate (**r**), succinate (**s**) and aspartate (**t**) in mock and SLC7A11 overexpressing cells following 24 h of ^13^C-glucose labelling. Data plotted are mean ± s.d. of n=5 wells.

